# Platform of single-molecule enzyme activity-based liquid biopsy for detection of pancreatic adenocarcinoma at early stages

**DOI:** 10.1101/2025.08.07.669035

**Authors:** Shingo Sakamoto, Hideto Hiraide, Tadahaya Mizuno, Mayano Minoda, Norimichi Nagano, Misa Suzuki, Nozomi Iwakura, Ayumu Kashiro, Satoshi Nara, Chigusa Morizane, Susumu Hijioka, Kazufumi Honda, Yu Kagami, Rikiya Watanabe, Yasuteru Urano, Toru Komatsu

## Abstract

Liquid biopsy analyzes biomolecules in biofluids to provide critical information for the early detection of diseases and the development of more effective treatment strategies. Among these biomolecules, the functional states of proteins offer particularly valuable insights due to their direct correlation with phenotypic changes. However, current approaches for assessing protein function in liquid biopsies are often limited by low detection sensitivity. Here, we present a liquid biopsy platform that analyzes single-molecule protease/peptidase activity to detect pathological alterations in enzyme function in blood samples. The platform demonstrates potential for identifying patients with early-stage (stage I-II) pancreatic ductal adenocarcinoma (PDAC).

## Introduction

Liquid biopsy is a methodology for analyzing disease-related biomolecules to facilitate early detection and enhanced understanding of various diseases. It is especially useful for the early detection of various types of cancer, and a variety of liquid biopsy platforms are being developed for this purpose^1^. While many existing platforms target nucleic acids, methods that enable the analysis of protein functions could provide unique information directly related to disease-related phenotypic changes^2^. Here, we report a platform for analyzing single-molecule protease/peptidase activities using multi-color fluorogenic probes that enable detection of pathological changes in enzyme activities at proteoform levels. The proposed liquid biopsy platform can detect unique molecular signature patterns of blood samples of patients with pancreatic adenocarcinoma (PDAC) in the early stages (stages I-II), demonstrating the platform’s potential for detecting diseases based on changes in protein function.

### Development of a single-molecule platform for analyzing protease/peptidase activities in blood samples

Single-molecule enzyme activity was analyzed using a micro-fabricated chamber device. Enzyme solution is first loaded into the device, which contains 10^4^-10^5^ chambers with a volume in the femtoliter range^3^. The enzyme concentration is set such that 0 or 1 molecule of the target enzyme is theoretically loaded into each chamber, and the activity of individual molecules is analyzed. We have recently proposed the concept of single-molecule enzyme activity profile (SEAP) analysis^4^ that utilize multi-colored enzyme substrates to discriminate different enzyme molecules based on different reactivities toward individual substrates. The assay is especially powerful in discriminating the activities of different enzyme proteoforms arising from variations in enzyme subtypes, posttranslational modifications (PTMs), protein-protein interactions (PPIs), and folding states (**Figure 1a**)^5^.

**Figure 1.**
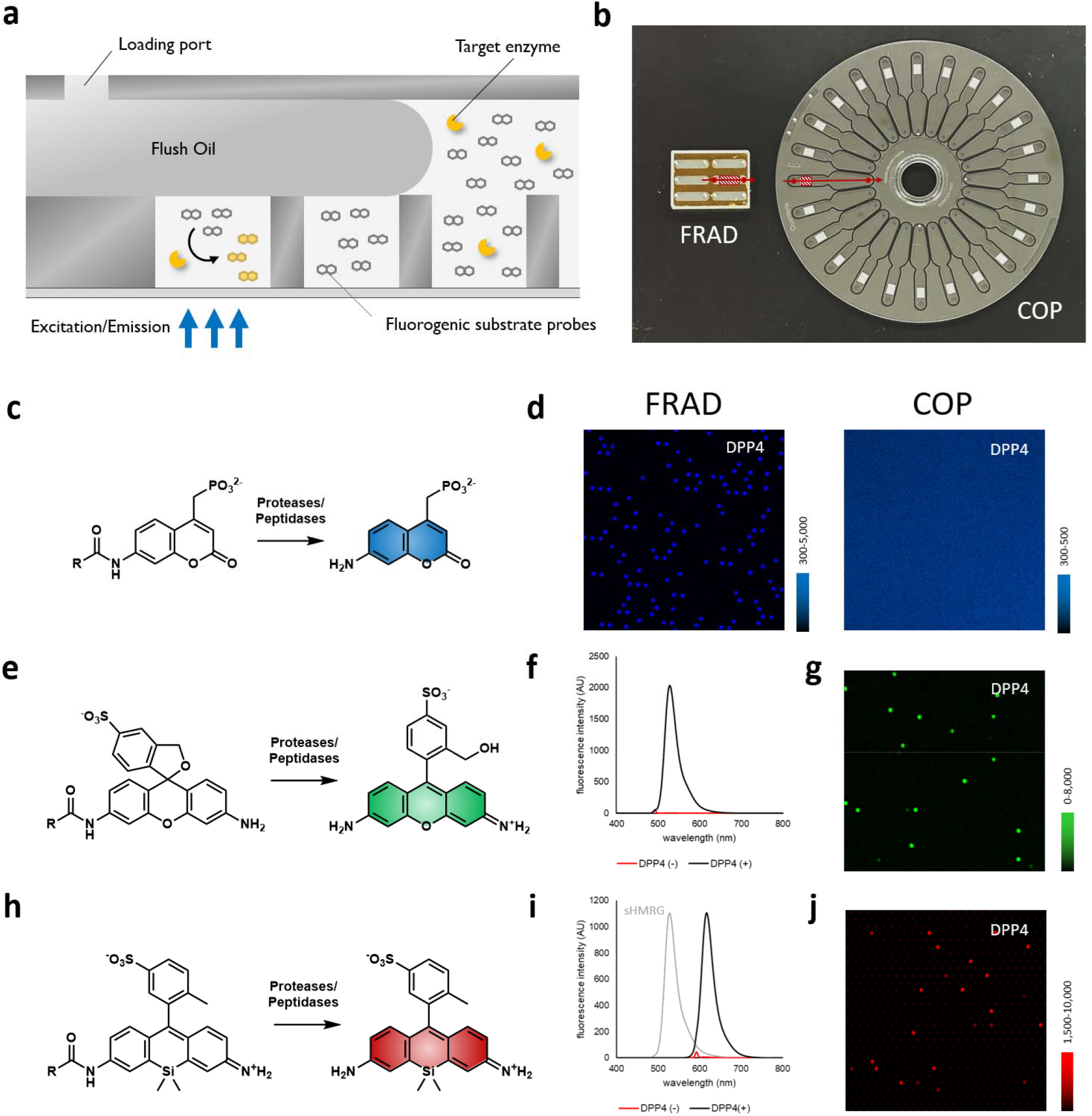
Development of multi-color fluorophores applicable to monitor protease/peptidase activities at the single-molecule level. (a) Concept of the single-molecule enzyme activity assay. (b) Images of the representative FRAD (left) and COP microdevice (right) used in the study. Arrows indicate the direction of solution flow, whereas circles indicate the loading and exit ports, and lines denote microchambers. (c) Structure of PMAC-based protease/peptidase probes. (d) Fluorescence images of FRAD and COP microdevice loaded with EP-PMAC (100 μM), blood samples of healthy subjects (1/20,000 for FRAD and 1/5,000 for COP microdevice) in HEPES-Na buffer (100 mM, pH 7.4) containing CaCl_2_ (1 mM), MgCl_2_ (1 mM), DTT (100 μM), and Triton X-100 (3 mM for FRAD and 250 μM for COP microdevice), incubated at 25°C for 4 h. (e) Structure of sHMRG-based protease/peptidase probes. (f) Fluorescence spectra of GP-sHMRG (3.3 μM) before and after mixing with recombinant DPP4 (10 ng/mL) in HEPES-Na buffer (100 mM, pH 7.4) and incubation for 30 min. λ_ex_ = 490 nm. (g) Fluorescence images of COP microdevice loaded with GP-sHMRG (30 μM), blood samples of healthy subjects (1/5,000) in HEPES-Na buffer (100 mM, pH 7.4) containing CaCl_2_ (1 mM), MgCl_2_ (1 mM), DTT (100 μM), Triton X-100 (250 μM), incubated at 25°C for 2 h. (h) Structure of sSiR-based protease/peptidase probes. (i) Fluorescence spectra of EP-sSiR (3.3 μM) before and after mixing with recombinant DPP4 (10 ng/mL) in HEPES-Na buffer (100 mM, pH 7.4) and incubation for 90 min. λ_ex_ = 490 nm. (j) Fluorescence images of COP microdevice loaded with EP-sSiR (30 μM), blood samples of healthy subjects (1/5,000) in HEPES-Na buffer (100 mM, pH 7.4) containing CaCl_2_ (1 mM), MgCl_2_ (1 mM), DTT (100 μM), Triton X-100 (250 μM), incubated at 25°C for 2 h.

Proteases/peptidases represent a major class of enzymes exhibiting a range of biological functions, including amino acid metabolism, signal transduction, extracellular matrix remodeling, and cell death^6,7^. Several studies have described changes in protease/peptidase activities in various diseases, revealing the potential of these enzymes as a promising foundation for the development of activity-based biomarkers^8–10^. Although a single-molecule protease/peptidase activity assay employing a coumarin fluorophore has been developed^10^, this method has three major limitations. First, the weak fluorescence of the coumarin dye results in low detection sensitivity, making it difficult to robustly quantify enzyme activity across a range of targets^11^. Second, the low penetration of UV excitation/emission light restricts the choice of microdevices; reliable signal detection was only possible with glass-bottomed devices such as femtoliter droplet array device (FRAD)^12^, and not with mass-producible polymer-based devices such as those made of cyclic olefin polymer (COP) (**Figure 1b-d**)^13,14^. Third, the assay was limited to single-channel (single-color) analysis, which restricted the ability to capture subtle differences in enzyme activity and complex formation that could be resolved through multi-channel (multi-color) analysis^4^.

To overcome the limitations, we designed a set of visible-light excitable fluorogenic probe scaffolds, sulfonated hydroxymethylrhodamine green (sHMRG, for green fluorescent probe) and sulfonated Si-rhodamine (sSiR, for red fluorescent probe), suitable for detecting single-molecule protease/peptidase activities with multi-colored SEAP analysis^4^. We recently synthesized fluorogenic probe to detect NAD(P)H^11^ using sHMRG as a fluorophore. As this fluorophore exhibits longer excitation/emission and higher ε × Φ_FL_ compared with coumarins, we hypothesized that we could use this scaffold to design green fluorescent protease/peptidase probes (**Figure 1e**). We established a scheme to efficiently synthesize sHMRG and tested peptide-modified probes for detecting protease/peptidase activity in a microdevice (**Scheme S1**). We initially designed and synthesized a DPP4 substrate by attaching a Gly-Pro (GP) sequence to sHMRG^15^. The resulting GP-sHMRG reacted with DPP4 to produce a >600-fold increase in fluorescence (**Figures 1f, S1, S2**). As expected, the probe was able to report single-molecule DPP4 activity in both FRAD and COP-based microdevices (**Figures 1g, S3)**. Besides widening the choice of microdevice, the brightness of the fluorophore was also beneficial for the rapid detection of the target. In detection of DPP4 activity in FRAD, GP-sHMRG gave the deired signal within 5 min, whereas the conventional coumarin-based probe EP-PMAC required at least 45 min for detection of reliable signals (**Figure S4**). In another case, increased brightness of the fluorophore provides a significant advantage in detection of enzymes with relatively weak single-molecule activity. Acylamino acid releasing enzyme (APEH) cleaves N-acylated amino acids such as Ac-Ala, Ac-Met, and fMet^16^. Single-molecule activity of APEH could not be detected using the coumarin-based probe (Ac-Ala-PMAC) with either a FRAD or COP-based device, but use of the sHMRG-based probe Ac-Ala-sHMRG led to successful detection of APEH activity with both microdevices (**Figure S5**).

In the same manner, we developed a red fluorescent fluorophore sSiR by modifying red-emitting Si-rhodamine^17,18^ with sulfonic acid (**Figure 1h, Scheme S2**). After attachment of substrate peptides for DPP4, the fluorophore reports the enzyme activity through a shift in the absorbance and fluorescence wavelength (**Figures 1i, S6, S2**). Due to its high hydrophilicity from sulfonic acid substitution, the activity of single-molecule DPP4 could be clearly monitored in microdevice-based assays as well (**Figure 1j**).

### Detection of single-molecule activity of DPP4 and CD13 in blood samples

Using the developed multi-color, single-molecule peptidase activity assay, two enzymes, CD13 and DPP4, were chosen as targets for activity monitoring in blood samples, as altered CD13 and DPP4 activity were observed in blood samples from PDAC (stage I-II) patients in our initial screening^10^ (**Table S1**). A previous study detected high– and low–DPP4 activity species in blood at the single-molecule level. The underlying molecular mechanisms of these species were unclear based on data from the original single-color assays^10^, but the two-colored single-molecule assay revealed that the varying activity derived from a homodimer of DPP4 (high DPP4 activity) and heterodimer of DPP4 with fibroblast activating protein α (FAPα)^19^. DPP4 and FAPα are both prolyl endopeptidases. DPP4 exhibits dipeptidyl peptidase–like activity, whereas FAPα prefers substrates with a masked *N*-terminus^20,21^. Using a combination of Ac-GP-sHMRG (green, targeting FAPα) and EP-sSiR (red, targeting DPP4), most spots with the half activity of dimeric DPP4 exhibited FAPα activity, indicating they are derived from a heterodimer of DPP4 with FAPα (**Figures 2a, 2b, S7**). Although studies on DPP4–FAPα are currently limited^19^, we found that the DPP4–FAPα heterodimer does not undergo dissociation/association equilibrium with the DPP4 or FAPα homodimers over a timescale of hours (**Figure S8**) Therefore, the presence of this molecular species in the blood is likely to reflect its formation in the cells/tissue of origin. A comparison of samples from PDAC patients and healthy subjects indicated a decrease in DPP4-FAPα heterodimer in blood samples of PDAC patients.

**Figure 2.**
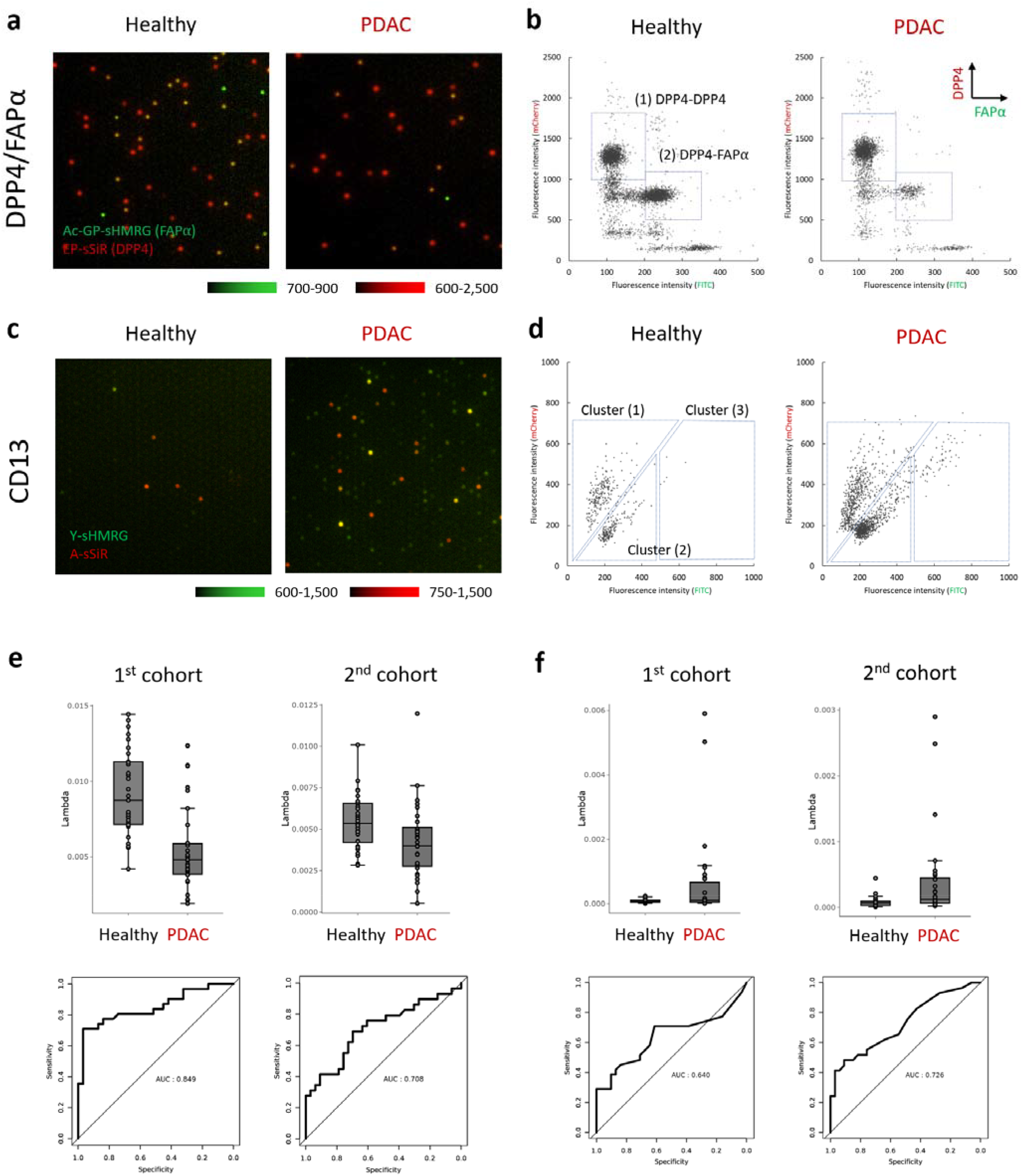
Multi-color single-molecule protease/peptidase assay to acquire the pathological activity patterns of enzymes in patients with pancreatic tumors. (a) Fluorescence images of microwells after loading Ac-GP-sHMRG (green, FAPα substrate, 30 μM), EP-sSiR (red, DPP4 substrate, 30 μM), IRDye800 (10 μM), and blood samples from healthy human subjects or PDAC patients (1/5,000 dilution) in HEPES-Na buffer (100 mM, pH 7.4) containing CaCl_2_ (1 mM), MgCl_2_ (1 mM), DTT (100 μM), Triton X-100 (150 μM), and bovine serum albumin (BSA; 1% w/v), incubated at 25°C for 4 h. (b) Scattered plot of (a). Horizontal axis indicates Ac-GP-sHMRG, and vertical axis indicates EP-sSiR. (c) Fluorescence images of microwells after loading Y-sHMRG (green, 30 μM), A-sSiR, (red, 30 μM), IRDye800 (10 μM), and blood samples from healthy human subjects or PDAC patients (1/2,000 dilution) in HEPES-Na buffer (100 mM, pH 7.4), CaCl_2_ (1 mM), MgCl_2_ (1 mM), DTT (100 μM), Triton X-100 (150 μM), and BSA (1% w/v), incubated at 25°C for 2 h. (d) Scattered plot of (c). Horizontal axis indicates Y-sHMRG, and vertical axis indicates A-sSiR. (e) Dot plot of lambda and receiver operating characteristic (ROC) curves of DPP4-FAPα heterodimers in healthy subjects and PDAC patients in 1^st^ and 2^nd^ cohort. (f) Dot plot of lambda and ROC curves of high-activity CD13 species in healthy subjects and PDAC patients in 1^st^ and 2^nd^ cohort.

For analysis of CD13 activity, the Tyr-sHMRG (Y-sHMRG)/Ala-sSiR (A-SiR) probe pair was tested (**Figures S9, S10**). This analysis revealed three major reactivity clusters in blood samples: (1) a cluster responding to both Y-sHMRG and A-sSiR, (2) a cluster responding primarily to Y-sHMRG, which overlapped with that of recombinant CD13, and (3) a cluster exhibiting higher activity than recombinant CD13 (**Figure 2c, 2d**). The substrate preferences of clusters (2) and (3) were similar, so cluster (3) seemed to reflect the molecular species consisting of multiple CD13 molecules. In samples from PDAC patients, the robust change was observed in cluster (3), which was almost undetectable in blood samples of healthy subjects.

### Automation of the assay and evaluation of diagnostic potential of biomarkers

Next, we analyzed multiple blood samples to test the feasibility of the biomarker candidates for the diagnosis of PDAC in the early stages (stages I-II). As we were able to use the mass-producible COP microdevice, the loading and fluorescence detection processes can be standardized and automated for increased throughput and reliability. To automate loading, we employed an automated siMoA digital ELISA system^14^. The experimental protocol was designed to mix the contents of 96-well plate wells (blood samples) with the solution in the reagent bottle (probe), load the mixed solution into a COP-based microdevice, and seal the device with flush oil using the SiMoA system (**Figure S11**). Fluorescence images of the COP-based microdevice were then acquired using an epifluorescence microscope. With this protocol, the loading of 24 samples was completed within 30 min, and clear signals were detected after incubation for a few hours (**Figure S12**). The overall process exhibited high reliability and repeatability, with coefficient of variation (CV) of most parameters remained less than 10% (**Figures S13, S14**). Next, multiple blood samples from PDAC patients and healthy subjects were analyzed (**Figures 2e, 2f, S15, S16**). Samples from 31 patients with PDAC (stages I-II) and 31 healthy subjects were analyzed (**Table S1**), resulting in area under curve (AUC) values of 0.85 for DPP4-FAPα and 0.64 for high-activity CD13. It is notable that these results were acquired for patients with PDAC in the early stages (stages I-II), for which the currently used biomarker CA19-9 is insufficient^22,23^.

Although this study was not designed as a prospective cohort analysis, we sought to ensure the reliability of our findings by validating the results using an independent cohort consisting of 33 healthy controls and 29 PDAC patients (stage I-II) collected at a different medical institution (**Table S1**). The reproducibility of detecting DPP4 and CD13 activity changes across independent cohorts underscores the robustness of these markers for detection of early stage PDAC. The same analysis protocols were applied to the 2^nd^ cohort, resulting in AUC values of 0.71 for DPP4 and 0.73 for CD13 (**Figures 2e, 2f, S15, S16**). Although the AUC values for the discovery (1^st^) and validation (2^nd^) cohorts were generally consistent, minor fluctuations were observed. Given the sample size of approximately 30 individuals per group, variability was expected due to individual differences, such as from healthy subjects with marginally elevated enzymatic activity. Nevertheless, these differences remained within a reasonable range, supporting the overall robustness of the markers.

### Deep learning–based extraction of molecular signature patterns in SEAP datasets

As described above, we were able to characterize disease-related changes in the activity of specific enzymes, focusing on one enzyme cluster each for DPP4 and CD13. However, both assays detected multiple clusters, for which changes in the number of constituents, activity intensity, or distribution could reveal biologically meaningful information^5^. To enable unbiased extraction of latent features from SEAP datasets, we developed an unsupervised representation learning-based analysis platform that leverages neural networks trained on multidimensional histograms (**Figure 3a**). A key challenge in biomarker discovery is extracting informative signatures from relatively small sample sizes (typically <100 samples). Therefore, we incorporated a variational autoencoder (VAE)–based pretraining framework to encompass the overall features of enzyme activities across all samples. The model was pretrained using combined data from both the 1^st^ and 2^nd^ cohorts, which enabled the unsupervised extraction of latent features without label information for downstream sample classification. The learning curves (**Figure 3b**) exhibited smooth and parallel decreases in both training and validation losses over epochs, suggesting stable optimization and effective representation learning, without signs of overfitting. The features are presented using 2-dimensional embedding, and unique clusters of PDAC patients were observed for both DPP4 and CD13 (**Figure 3c**). By showing the 3 most similar and least similar samples from a randomly selected samples, it appeared that the system seemed to capture features observed uniquely in PDAC (**Figures 3d, S17, S18**); for DPP4, samples with weak DPP4-FAPα heterodimers were read out to be similar (**Figure S17**), and for CD13, samples with an increased number of high-activity CD13 species were read out to be similar (**Figure S18**). These results highlight the model’s capacity for detecting biologically interpretable variations without manual thresholding. We then applied this analytical framework to assess the combined molecular signatures of DPP4 and CD13. The disease discrimination model was constructed by training the model on 1^st^ cohort and testing it on 2^nd^ cohort. It gave a classification performance characterized by an area under the receiver operating characteristic curve (AUROC) value of 0.77 ± 0.01 (n = 3, mean ± SD) and accuracy of 0.75 ± 0.01 (n = 3, mean ± SD) for distinguishing stage I-II PDAC samples from healthy control samples (**Figure 3e**). The value was in good agreement with the AUC values of threshold-based analysis, supporting the notion that the feature extraction model captures reproducible patterns across independent cohorts. Revisiting the 2-dimensional embedding (**Figure 3c**), multiple clusters were generated, but the tumor-specific cluster primarily included patients exhibiting low DPP4-FAPα heterodimer level and/or high CD13 activity. Other molecular features might not differentiate PDAC patients from healthy subjects but could reflect other health conditions or individual differences.

**Figure 3.**
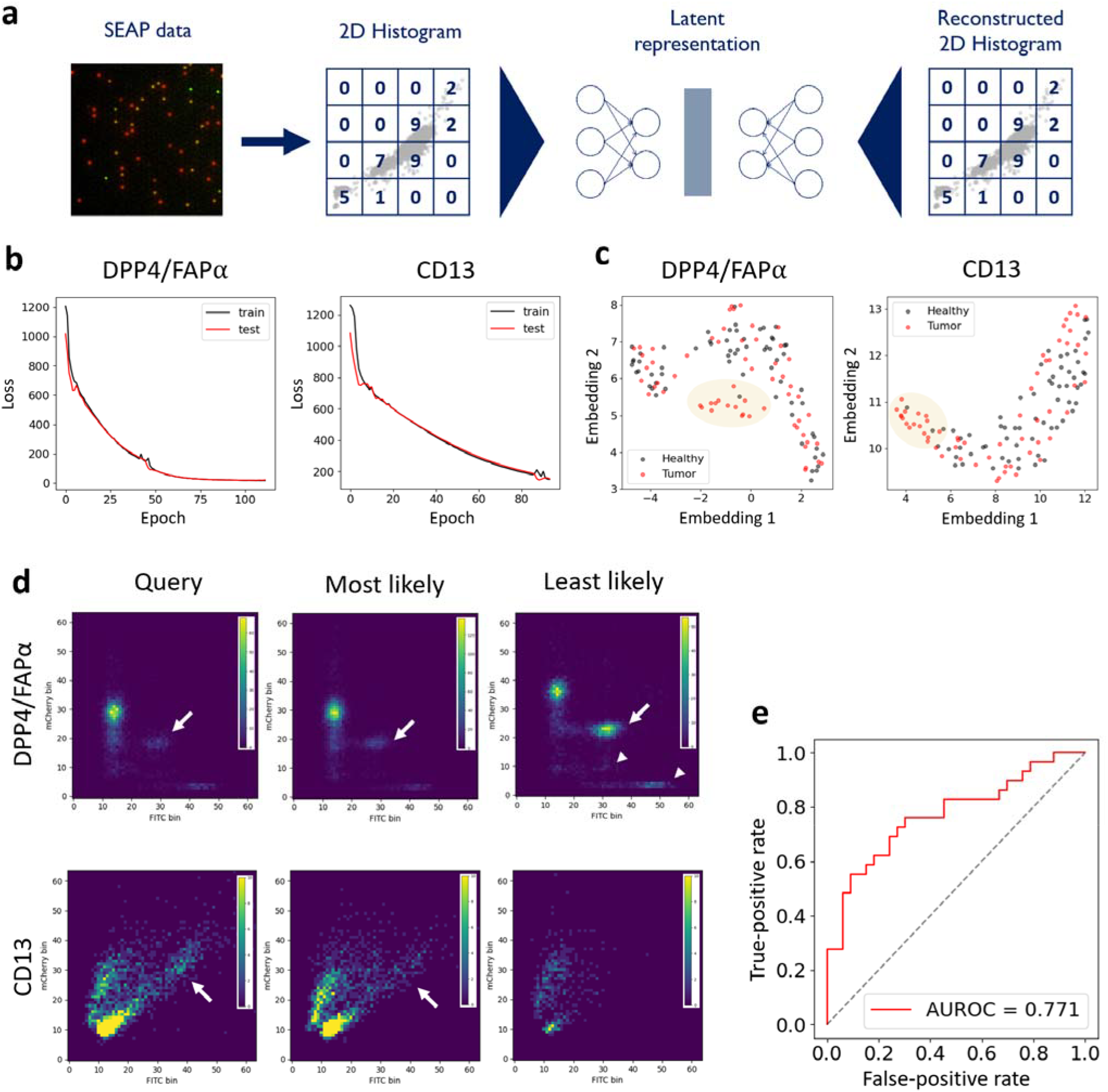
(a) Schematic view of data encoding and analysis. (b) Learning curves for loss per each query in train and test studies for DPP4 and CD13. (c) Latent space representation of multiple features in the VAE study. Clusters baring tumor-specific molecular signatures (low DPP4-FAPα and high CD13) are indicated as pale yellow circles. (d) Representative examples of most and least likely specimens for each query sample for the analysis of DPP4 and CD13. PDAC-specific changes are shown as white arrows, and other signature patterns considered to contribute to likeliness are indicated as white arrowheads. More data are shown in **Figures S17** and **S18**. (e) AUROC curves for discriminating between healthy subjects and early stage (stages I-II) PDAC patients, trained on 1^st^ cohort and evaluated on 2^nd^ cohort.

## Discussion

PDAC remains one of the leading causes of cancer-related mortality, largely because it is often diagnosed at advanced stages, when curative treatment is rarely possible^24,25^. However, early detection through minimally invasive liquid biopsy holds promise for improving patient outcomes^22^. Here, we report a novel liquid biopsy platform based on measuring the activity of proteases in plasma at the single-molecule level. Among the various modalities used in liquid biopsy, the analysis of protein function, changes in which are directly connected to phenotypic changes, lies downstream of the central dogma^2^. The advantages of protein activity analysis can be summarized by the concept of activity-based diagnostics^26^, but current methodologies involving bulk analysis of 10^6^-10^9^ molecules as a bundle suffer from low detection sensitivity and an inability to discriminate between different proteoforms, molecular forms resulting from PTMs and PPIs^27^. Methods enabling single-molecule enzyme activity analysis could enable the discrimination of various proteoforms simultaneously^5^, and we therefore developed an assay platform for detecting the activity of PDAC-related enzymes.

The pancreas is a source tissue for a variety of proteases/peptidases. Therefore, we focused our biomarker research on identifying useful proteases/peptidases. In this study, we characterized changes in the activity of DPP4-FAPα and CD13 proteoforms in blood samples collected from PDAC patients (**Figure 2e, 2f**). Both DPP4 and FAPα are generated as membrane-bound proteins, but molecules detected in the plasma are thought to reflect the soluble forms of these enzymes cleaved from membranes by various proteases^28^. The kidney is a major DPP4-expressing organ, but circulating soluble DPP4 does not originate from the kidney but rather from immune cells, fibroblasts, and endothelial cells^28^. FAPα is generally absent in most normal adult tissues, and the cellular sources contributing to its presence in the circulation remain poorly characterized^29^. With regard to the origin of DPP4-FAPα heterodimers, it did not occur from equilibrium between dissociation/association (**Figure S8**), so changes in the blood reflect the altered generation or secretion of DPP4-FAPα by source cells/tissues. Pancreatic α-cells represent a potential source, as they are known to express both DPP4 and FAPα^30^. Another possible source is tumor-associated fibroblasts^31^. In either case, it seems that the remodeled stromal microenvironment in PDAC contributes to altered secretion of DPP4-FAPα into the bloodstream.

The observed elevation in total CD13 activity was consistent with a previous study that identified CD13 as a candidate biomarker for PDAC^10^, but results of the multi-color assay of the present study revealed that multiple active species are present in the bloodstream. This observation in turn led to the identification of a high-activity CD13 species that could serve as a reliable biomarker of PDAC, as this species is rarely detected in healthy subjects. The high-activity species is considered to consist of 2-3 CD13 molecules corresponding to recombinant CD13 (**Figure S10**). The distribution cannot be explained from simple Poisson’s model^32^, indicating that there are molecular species consisting of multiple CD13 molecules. The previous study indicated that CD13 associates with high density lipoprotein (HDL) particles in blood samples of PDAC patients^10^, raising the possibility that the source of this species is CD13 interacting with HDL; however, further studies are required to verify this possibility and elucidate the underlying molecular mechanism.

The key message of this study is that the platform offers both high stability and rapid measurement performance, making it readily applicable as a liquid biopsy tool. Its ability to reliably detect disease-related changes in molecular signature patterns highlights its potential for early-stage PDAC detection. In this study, we also developed an unbiased data analysis platform using a deep learning–based analytical framework. Data-driven approaches have proven effective in flow cytometry^33^ and fluorescence imaging^34^ for extracting subtle, high-dimensional differences between biological samples. Similar advantages can be expected in SEAP analysis as well, particularly when applied to larger datasets in future studies. The platform is inherently scalable to higher-dimensional data, broader enzyme panels, and larger patient cohorts, offering the potential to enhance diagnostic precision through systematic and unbiased molecular profiling.

One limitation of this study is the relatively small number of samples analyzed, owing to the limited availability of blood from patients with early-stage PDAC (stage I–II). While our VAE-based framework showed promise in capturing meaningful molecular signatures, the current sample size constrains a comprehensive evaluation of its ability to represent broader clinical diversity. Expanding the dataset will be essential to fully validate the clinical utility of the platform. Another important direction for future research is the careful expansion of the enzyme panel used in the assay. In both threshold-based and deep learning–based approaches, AUC values reached approximately 0.75–0.8, indicating that some early-stage PDAC patients may not exhibit significant alterations in the two enzymes currently analyzed. Incorporating one or two additional biomarker enzymes, selected to complement the existing ones, could help improve patient coverage and enhance diagnostic sensitivity. The present study establishes a foundation for extending this approach to a broader, yet focused, set of enzymes, particularly those relevant to gastroenterological diseases. Systematic screening of additional proteases and related enzyme classes will identify such complementary biomarkers and further improve diagnostic performance.

## Conclusion

In summary, we have developed a single-molecule enzyme activity assay capable of resolving distinct proteoforms in blood with high sensitivity and molecular specificity. Applying this platform to early stage PDAC, we identified enzymatically distinct species of DPP4/FAPα and CD13 that serve as promising biomarker candidates. This work not only provides proof-of-concept for functional proteoform-based diagnostics, it also opens a path toward scalable, activity-based screening of disease-relevant enzymes in clinical samples. Beyond its application to blood-based cancer diagnostics, our single-molecule enzyme activity assay offers a generalizable strategy for functionally resolving proteoforms in complex biological fluids.

## Author Contributions

T. K. conceived the experimental design. S. S., M. M., and T. K., synthesized and characterized compounds. S. S., H. H., M. M., N. N., M. S., N. I., and T. K. acquired the experimental data. T. M. constructed the data analysis platform. A. K., S. N., C. M., S. H. and K.H. prepared plasma samples. R. W. prepared microdevice. The experimental data were analyzed under the supervision of T. M., Y. K., K. H., R. W., Y. U. and T. K. The manuscript was written by T. K.

## Acknowledgements

This work was financially supported by MEXT (20H04694, 21A303, 22H02217, 23K23484, 25K01911 and 25K22520 to T. K., 21K06663 to T. M., 25H00241 for N. N.), JST (PRESTO (13414915), PRESTO Network (17949814) and FOREST (24012649) to T. K., CREST (19204926) to R. W. and T. K., START (20353017) to T. M., K. H., R. W., and T. K.), and AMED (FORCE (22581634) to T. M., K. H., R. W., and T. K., P-PROMOTE (25131640) to T. M., K. H., and T. K., P-PROMOTE (18cm0106403h0003) to K. H., P-CREATE (25ama221431h0002) to K. H. T. K. received support from the Naito Foundation, The Mochida Memorial Foundation for Medical and Pharmaceutical Research, Chugai Foundation for Innovative Drug Discovery Science, MSD Life Science Foundation, Hoansha Foundation, and University of Tokyo Gap Fund Program. We thank Dr. Oketani for collecting the plasma samples.

## REFERENCES

(1) Ignatiadis, M.; Sledge, G. W.; Jeffrey, S. S. Liquid Biopsy Enters the Clinic - Implementation Issues and Future Challenges. Nat. Rev. Clin. Oncol. 2021, 18 (5), 297–312. 10.1038/s41571-020-00457-x.

(2) Saghatelian, A.; Cravatt, B. F. Assignment of Protein Function in the Postgenomic Era. Nat. Chem. Biol. 2005, 1 (3), 129. 10.1038/nchembio0805-130.

(3) Noji, H.; Minagawa, Y.; Ueno, H. Enzyme-Based Digital Bioassay Technology - Key Strategies and Future Perspectives. Lab Chip 2022, 22 (17), 3092–3109. 10.1039/d2lc00223j.

(4) Sakamoto, S.; Komatsu, T.; Watanabe, R.; Zhang, Y.; Inoue, T.; Kawaguchi, M.; Nakagawa, H.; Ueno, T.; Okusaka, T.; Honda, K.; Noji, H.; Urano, Y. Multiplexed Single-Molecule Enzyme Activity Analysis for Counting Disease-Related Proteins in Biological Samples. Sci. Adv. 2020, 6 (11), eaay0888. 10.1126/sciadv.aay0888.

(5) Komatsu, T.; Mizuno, T. Single-Molecule Enzyme Activity Analysis for Illuminating Pathological Proteoforms. ACS Cent. Sci. 2025, 11 (7), 1041–1051. 10.1021/acscentsci.5c00100.

(6) López-Otín, C.; Bond, J. S. Proteases: Multifunctional Enzymes in Life and Disease. J. Biol. Chem. 2008, 283 (45), 30433–30437. 10.1074/jbc.R800035200.

(7) Long, J. Z.; Cravatt, B. F. The Metabolic Serine Hydrolases and Their Functions in Mammalian Physiology and Disease. Chem. Rev. 2011, 111 (10), 6022–6063. 10.1021/cr200075y.

(8) Onagi, J.; Komatsu, T.; Ichihashi, Y.; Kuriki, Y.; Kamiya, M.; Terai, T.; Ueno, T.; Hanaoka, K.; Matsuzaki, H.; Hata, K.; Watanabe, T.; Nagano, T.; Urano, Y. Discovery of Cell-Type-Specific and Disease-Related Enzymatic Activity Changes via Global Evaluation of Peptide Metabolism. J. Am. Chem. Soc. 2017, 139 (9), 3465–3472. 10.1021/jacs.6b11376.

(9) Kuriki, Y.; Yoshioka, T.; Kamiya, M.; Komatsu, T.; Takamaru, H.; Fujita, K.; Iwaki, H.; Nanjo, A.; Akagi, Y.; Takeshita, K.; Hino, H.; Hino, R.; Kojima, R.; Ueno, T.; Hanaoka, K.; Abe, S.; Saito, Y.; Nakajima, J.; Urano, Y. Development of a Fluorescent Probe Library Enabling Efficient Screening of Tumour-Imaging Probes Based on Discovery of Biomarker Enzymatic Activities. Chem. Sci. 2022, 13, 4474–4481. 10.1039/D1SC06889J.

(10) Sakamoto, S.; Hiraide, H.; Minoda, M.; Iwakura, N.; Suzuki, M.; Ando, J.; Takahashi, C.; Takahashi, I.; Murai, K.; Kagami, Y.; Mizuno, T.; Koike, T.; Nara, S.; Morizane, C.; Hijioka, S.; Kashiro, A.; Honda, K.; Watanabe, R.; Urano, Y.; Komatsu, T. Identification of Activity-Based Biomarkers for Early-Stage Pancreatic Tumors in Blood Using Single-Molecule Enzyme Activity Screening. Cell Rep. Methods 2024, 4, 100688. 10.1016/j.crmeth.2023.100688.

(11) Minoda, M.; Hatakeyama, J.; Nagano, N.; Mizuno, T.; Iwasaka, T.; Shiga, S.; Takahashi, K.; Hiraide, H.; Sakamoto, S.; Kagami, Y.; Kashiro, A.; Honda, K.; Sugiura, Y.; Mishima, K.; Mishima, M. K.; Kusuhara, H.; Urano, Y.; Komatsu, T. Single-Molecule Oxidoreductase Activity Analysis for Activity-Based Diagnosis Based on Proteoform Alterations. J. Am. Chem. Soc. 2025, 147 (6), 4743–4751. 10.1021/jacs.4c07624.

(12) Sakakihara, S.; Araki, S.; Iino, R.; Noji, H. A Single-Molecule Enzymatic Assay in a Directly Accessible Femtoliter Droplet Array. Lab Chip 2010, 10 (24), 3355–3362. 10.1039/c0lc00062k.

(13) Zhang, H.; Nie, S.; Etson, C. M.; Wang, R. M.; Walt, D. R. Oil-Sealed Femtoliter Fiber-Optic Arrays for Single Molecule Analysis. Lab Chip 2012, 12 (12), 2229–2239. 10.1039/c2lc21113k.

(14) Kan, C. W.; Rivnak, A. J.; Campbell, T. G.; Piech, T.; Rissin, D. M.; Mösl, M.; Petera, A.; Niederberger, H. P.; Minnehan, K. A.; Patel, P. P.; Ferrell, E. P.; Meyer, R. E.; Chang, L.; Wilson, D. H.; Fournier, D. R.; Duffy, D. C. Isolation and Detection of Single Molecules on Paramagnetic Beads Using Sequential Fluid Flows in Microfabricated Polymer Array Assemblies. Lab Chip 2012, 12 (5), 977–985. 10.1039/c2lc20744c.

(15) Onoyama, H.; Kamiya, M.; Kuriki, Y.; Komatsu, T.; Abe, H.; Tsuji, Y.; Yagi, K.; Yamagata, Y.; Aikou, S.; Nishida, M.; Mori, K.; Yamashita, H.; Fujishiro, M.; Nomura, S.; Shimizu, N.; Fukayama, M.; Koike, K.; Urano, Y.; Seto, Y. Rapid and Sensitive Detection of Early Esophageal Squamous Cell Carcinoma with Fluorescence Probe Targeting Dipeptidylpeptidase IV. Sci. Rep. 2016, 6 (1), 26399. 10.1038/srep26399.

(16) Komatsu, T.; Hanaoka, K.; Adibekian, A.; Yoshioka, K.; Terai, T.; Ueno, T.; Kawaguchi, M.; Cravatt, B.; Nagano, T. Diced Electrophoresis Gel Assay for Screening Enzymes with Specified Activities. J. Am. Chem. Soc. 2013, 135 (16), 6002–6005. 10.1021/ja401792d.

(17) Kushida, Y.; Nagano, T.; Hanaoka, K. Silicon-Substituted Xanthene Dyes and Their Applications in Bioimaging. Analyst. Royal Society of Chemistry February 7, 2015, pp 685–695. 10.1039/c4an01172d.

(18) Kushida, Y.; Hanaoka, K.; Komatsu, T.; Terai, T.; Ueno, T.; Yoshida, K.; Uchiyama, M.; Nagano, T. Red Fluorescent Scaffold for Highly Sensitive Protease Activity Probes. Bioorg. Med. Chem. Lett. 2012, 22 (12), 3908–3911. 10.1016/j.bmcl.2012.04.114.

(19) Scanlan, M. J.; Raj, B. K. M.; Calvo, B.; Garin-Chesa, P.; Pilar Sanz-Moncasi, M.; Healeyt, J. H.; Old, L. J.; Rerrfig, W. J. Molecular Cloning of Fibroblast Activation Protein a, a Member of the Serine Protease Family Selectively Expressed in Stromal Fibroblasts of Epithelial Cancers (Type I Integral Membrane Protein/Wound Heaflng/Sarcoma/Dipeptdyl Peps IV/CD26). Proc. Natl. Acad. Sci. USA 1994, 91, 5657–5661.

(20) Sakabe, M.; Asanuma, D.; Kamiya, M.; Iwatate, R. J.; Hanaoka, K.; Terai, T.; Nagano, T.; Urano, Y. Rational Design of Highly Sensitive Fluorescence Probes for Protease and Glycosidase Based on Precisely Controlled Spirocyclization. J. Am. Chem. Soc. 2013, 135 (1), 409–414. 10.1021/ja309688m.

(21) Aertgeerts, K.; Levin, I.; Shi, L.; Snell, G. P.; Jennings, A.; Prasad, G. S.; Zhang, Y.; Kraus, M. L.; Salakian, S.; Sridhar, V.; Wijnands, R.; Tennant, M. G. Structural and Kinetic Analysis of the Substrate Specificity of Human Fibroblast Activation Protein α. J. Biol. Chem. 2005, 280 (20), 19441–19444. 10.1074/jbc.C500092200.

(22) Kato, S.; Honda, K. Use of Biomarkers and Imaging for Early Detection of Pancreatic Cancer. Cancers (Basel) 2020, 12 (7), 1–21. 10.3390/cancers12071965.

(23) Ferrone, C. R.; Finkelstein, D. M.; Thayer, S. P.; Muzikansky, A.; Fernandez-Del Castillo, C.; Warshaw, A. L. Perioperative CA19-9 Levels Can Predict Stage and Survival in Patients with Resectable Pancreatic Adenocarcinoma. J. Clin. Oncol. 2006, 24 (18), 2897–2902. 10.1200/JCO.2005.05.3934.

(24) Ryan, D. P.; Hong, T. S.; Bardeesy, N. Pancreatic Adenocarcinoma. N. Eng. J. Med. 2014, 371 (11), 1039–1049. 10.1056/NEJMra1404198.

(25) Siegel, R. L.; Miller, K. D.; Wagle, N. S.; Jemal, A. Cancer Statistics, 2023. CA Cancer J. Clin. 2023, 73 (1), 17–48. 10.3322/caac.21763.

(26) Soleimany, A. P.; Bhatia, S. N. Activity-Based Diagnostics: An Emerging Paradigm for Disease Detection and Monitoring. Trends Mol. Med. 2020, 26 (5), 450–468. 10.1016/j.molmed.2020.01.013.

(27) Aebersold, R.; Agar, J. N.; Amster, I. J.; Baker, M. S.; Bertozzi, C. R.; Boja, E. S.; Costello, C. E.; Cravatt, B. F.; Fenselau, C.; Garcia, B. A.; Ge, Y.; Gunawardena, J.; Hendrickson, R. C.; Hergenrother, P. J.; Huber, C. G.; Ivanov, A. R.; Jensen, O. N.; Jewett, M. C.; Kelleher, N. L.; Kiessling, L. L.; Krogan, N. J.; Larsen, M. R.; Loo, J. A.; Ogorzalek Loo, R. R.; Lundberg, E.; Maccoss, M. J.; Mallick, P.; Mootha, V. K.; Mrksich, M.; Muir, T. W.; Patrie, S. M.; Pesavento, J. J.; Pitteri, S. J.; Rodriguez, H.; Saghatelian, A.; Sandoval, W.; Schlüter, H.; Sechi, S.; Slavoff, S. A.; Smith, L. M.; Snyder, M. P.; Thomas, P. M.; Uhlén, M.; Van Eyk, J. E.; Vidal, M.; Walt, D. R.; White, F. M.; Williams, E. R.; Wohlschlager, T.; Wysocki, V. H.; Yates, N. A.; Young, N. L.; Zhang, B. How Many Human Proteoforms Are There? Nat. Chem. Biol. Nature Publishing Group February 14, 2018, pp 206–214. 10.1038/nchembio.2576.

(28) Wang, Z.; Grigo, C.; Steinbeck, J.; Von Hörsten, S.; Amann, K.; Daniel, C. Soluble DPP4 Originates in Part from Bone Marrow Cells and Not from the Kidney. Peptides (N.Y.) 2014, 57, 109–117. 10.1016/j.peptides.2014.05.006.

(29) Hamson, E. J.; Keane, F. M.; Tholen, S.; Schilling, O.; Gorrell, M. D. Understanding Fibroblast Activation Protein (FAP): Substrates, Activities, Expression and Targeting for Cancer Therapy. Proteomics Clin. Appl. 2014, 8 (5–6), 454–463. 10.1002/prca.201300095.

(30) Busek, P.; Hrabal, P.; Fric, P.; Sedo, A. Co-Expression of the Homologous Proteases Fibroblast Activation Protein and Dipeptidyl Peptidase-IV in the Adult Human Langerhans Islets. Histochem. Cell Biol. 2015, 143 (5), 497–504. 10.1007/s00418-014-1292-0.

(31) Lee, J.; Fassnacht, M.; Nair, S.; Boczkowski, D.; Gilboa, E. Tumor Immunotherapy Targeting Fibroblast Activation Protein, a Product Expressed in Tumor-Associated Fibroblasts. Cancer Res. 2005, 65 (23), 11156–11163. 10.1158/0008-5472.CAN-05-2805.

(32) Rissin, D. M.; Walt, D. R. Digital Concentration Readout of Single Enzyme Molecules Using Femtoliter Arrays and Poisson Statistics. Nano Lett. 2006, 6 (3), 520–523. 10.1021/nl060227d.

(33) Arvaniti, E.; Claassen, M. Sensitive Detection of Rare Disease-Associated Cell Subsets via Representation Learning. Nat. Commun. 2017, 8. 10.1038/ncomms14825.

(34) Chandrasekaran, S. N.; Ceulemans, H.; Boyd, J. D.; Carpenter, A. E. Image-Based Profiling for Drug Discovery: Due for a Machine-Learning Upgrade? Nat. Rev. Drug Discov. Nature Research February 1, 2021, pp 145–159. 10.1038/s41573-020-00117-w.

