## Supplementary_Information for "Platform of single-molecule enzyme activity-based liquid biopsy for detection of pancreatic adenocarcinoma at early stages"

**Contents**

**Methods**

**Supplementary methods for data encoding**

**Supplementary data and comments**

**Supplementary data for synthesis and characterization of compounds**

**Supplementary references**

### Methods

#### Materials

Reagents and solvents were of the best grade available, supplied by Tokyo Chemical Industries, Wako Pure Chemical, Sigma-Aldrich, Dojindo, Kanto Chemical Co., Watanabe Chemical Industries, and were used without further purification. Enzymes were purchased from Sigma-Aldrich and was used without further purification.

#### Enzymes

Recombinant DPP4 was purchased from R&D Biosystems (catalog # 9168-SE). Recombinant CD13 was purchased from R&D Biosystems (catalog #3815-ZN). Recombinant FAPα was purchased from R&D Biosystems (catalog #3715-SE).

#### Instruments

NMR spectra were recorded on a JEOL JNM-LA400 instrument at 400 MHz for $^{1}H$ NMR and at 100 MHz for $^{13}C$ NMR. Mass spectra (MS) were measured with a JEOL JMS-T100LC AccuToF (ESI). LC-MS analyses were performed on a Waters Acquity UPLC (H class)/QDa quadrupole MS analyzer or Acquity UPLC (H class)/Xevo TQD quadrupole MS/MS analyzer equipped with an Acquity UPLC BEH C18 column (Waters). Column chromatography using silica gel was performed on a MPLC system (Yamazen Smart Flash EPCLC AI-5805 (Tokyo, Japan)). Reversed-phase MPLC purification was performed on an Isolera One (Biotage) equipped with a SNAP Ultra C18 30 g (Biotage).

#### UV-Vis Absorption and fluorescence spectroscopy

UV-Visible spectra were obtained on a Shimadzu UV-1800. Fluorescence spectroscopic studies were performed on a Hitachi F7000. The slit width was 5 nm for both excitation and emission. The photomultiplier voltage was 400 V.

**Fluorescence microscopy**

Fluorescence images were acquired by fluorescence microscope (Ti2, Nikon) equipped with a 20× dry objective lens (Plan Apo 20×), sCMOS camera (ORCA-Fusion C14440, Hamamatsu Photonics), white LED illumination unit (X-Cite Xylis, Opto Science), moving stage and rotating stage. The stage was custom-made to hold Simoa disk (Quanterix) and rotate it using motorized rotation stage(OSMS-60YAW, Opto Sigma). For continuous monitoring of multiple device images, the rotating stage rotated the disk by 15° for each acquisition. Each image was acquired in tile scan mode of 3×4 (with overlap of 1%) with perfect focus. The assay was performed using a solution containing IR-dye 800 (10 μM) as the internal standard, and the focus was adjusted using its fluorescence. All operations were controlled by NIS-Element software (Nikon). The excitation and emission filters used were DAPI (mirror = 425 nm, ex. = 383-408 nm, Em. = 435-485 nm), FITC (mirror = 510 nm, Ex. = 460-500 nm, Em. = 510-560 nm), mCherry (mirror = 600 nm, ex. = 550-590 nm, Em. = 608-683 nm), and Cy7 (mirror = 743 nm, Ex. = 743 nm, Em. = 767 nm), respectively. Fluorescence signals of sHMRG-based probes were acquired with FITC filter and those of sSiR-based probes were acquired with mCherry filter.

**Single-molecule enzyme activity assay using SR-X**

The single-molecule enzyme activity assay was performed using the SR-X system (Quanterix Corp.). Biological samples were prepared in 96-well plates (Quanterix Corp.) at 2× concentration relative to the final concentration, whereas the probe solution was prepared at 2× concentration in a narrow-mouth bottle (Thermo Scientific). A 40 μL volume of probe solution and 40 μL of each biological sample were mixed and introduced into a Simoa Disc (Quanterix Corp.) using the SR-X system. Subsequently, Simoa Sealing Oil (Quanterix Corp.) was added to the Simoa Disc. After processing using the SR-X system, additional Simoa Sealing Oil was manually applied at the lane inlets of the Simoa Disc, and the lane outlets of the Simoa Disc were sealed with stickers to prevent evaporation of the reaction solution. The enzymatic activity in the chambers was measured using fluorescence microscopy. The images of 12 points were used for the analysis of DPP4, and 9 points were used for the analysis of CD13.

**Image processing**

Images were processed using the GA3 module of NIS Elements software (Nikon). First, all fluorescence images were background-corrected using a rolling ball correction (diameter = 3 μm). Then, ROIs were chosen by bright spot detection using DAPI and FITC filters (diameter = 3 μm), and irregular fluorescent spots derived from fluorescent debris or air bubbles were omitted by dilating the ROI (1.3 μm) and removing the overlapping ROIs by size (fill area < 40) and shape (circularity > 0.8) filter. The fluorescence signals were acquired as the mean of the signals from 3×3 pixels from the center of each ROI. Data were processed using Python, Excel, or Kaleidagraph software to construct activity histograms.

**Plasma samples and ethics statement**

Plasma samples were obtained from the Platform for Evaluating Biomarkers of Cancer Early Detection (P-EBED), with the Program for Promotion of Fundamental Studies in Health Sciences conducted by the National Institute of Biomedical Innovation of Japan, Health and Labour Sciences Research Grants from the Ministry of Health, Labor and Welfare of Japan, and P-CREATE of the Japan Agency for Medical Research and Development (AMED)[1–4]. Ethical approval for this study was obtained from the central ethics committees of Nippon Medical School (M-2021-002) and the ethical committee of Nippon Medical School (A-2020-032 and A-2020-044).

### Supplementary methods for data encoding

SEAP data were encoded according to the following processes. The model is available at https://github.com/mizuno-group/hist-vae.git.

#### 1. Data preprocessing

Data were acquired using the SEAP platform from two independent centers, referred to as cohort 1 and cohort 2. Each cohort comprised 62 samples, evenly split between healthy and cancerous tissue. For each specimen, data were collected as two-dimensional (2D) point clouds derived from dual-color imaging (FITC and mCherry), and data were collected separately for the surface markers DPP4 and CD13.

Prior to model input, value-based trimming was applied to filter out extreme values. For DPP4, the FITC channel was restricted to [0, 500] and the mCherry channel to [0, 2,500], whereas CD13 was trimmed to [0, 1,000] in both channels. After preprocessing, a total of 192,660 (DPP4) and 50,030 (CD13) points were obtained for cohort 1, and 176,167 (DPP4) and 50,851 (CD13) points were obtained for cohort 2.

#### 2. Data preprocessing and representation

SEAP-derived data consisted of unordered 2D point clouds representing spatial distributions of molecular signals. To enable convolutional neural network (CNN)–based learning, each point cloud was converted into a fixed-size 2D histogram. This was achieved by defining a uniform grid over the bounding box of each sample and binning the coordinates accordingly. Although the present study focused on 2D inputs, the pipeline supports extension to 1D and 3D data.

To enhance the robustness of the assay and facilitate self-supervised training, two independently subsampled versions of each point cloud were generated and subjected to light geometric transformations, including jittering, scaling, and translation. One of the resulting histograms was treated as a “clean” target and the other as “noisy” input for denoising reconstruction. Prior to model input, all histograms were log-transformed and normalized to [0,1] using group-specific maxima to ensure consistency across samples.

#### 3. Model architecture

A 2D convolutional variational autoencoder (VAE) was employed to learn latent representations of the histogram-encoded point clouds. The encoder consisted of residual convolutional blocks with down-sampling, whereas the decoder used transposed convolutional blocks with skip connections. The encoder mapped input histograms to the parameters of a multivariate Gaussian distribution in a latent space, characterized by its mean and log-variance. This distribution was sampled via the reparameterization trick. The decoder then reconstructed the original histogram from the sampled latent vector.

The total loss function, $\mathcal{L}$, was defined as a weighted sum of the reconstruction loss and the KL divergence, as follows:

$$\mathcal{L} = MSE(\hat{\boldsymbol{x}}, \boldsymbol{x}) + \beta \cdot D_{KL}\big(q(\boldsymbol{z}|\boldsymbol{x}) \parallel p(\boldsymbol{z})\big),$$

where $\hat{\boldsymbol{x}}$ and $\boldsymbol{x}$ denote the reconstructed and original histograms, $q(\boldsymbol{z}|\boldsymbol{x})$ represents the approximate posterior, $p(\boldsymbol{z}) \sim \mathcal{N}(\boldsymbol{0}, \boldsymbol{I})$ represents the prior, and $\beta$ is a tunable regularization coefficient.

**4. Representation learning**

The model was trained in a self-supervised manner to reconstruct clean histograms from noisy inputs. No class labels were used. Optimization was performed using the Adam optimizer with gradient clipping and gradient accumulation to ensure numerical stability during training. Training was conducted for a fixed number of epochs or until early stopping criteria were met based on the validation loss. Although a fine-tuning pipeline with classification heads was implemented, this pipeline was not utilized in the current study due to the limited dataset size.

**5. Implementation details**

All experiments were implemented in Python 3.10.12 using PyTorch 2.3.0 and executed on a single GPU machine (RTX3090, NVIDIA). GPU acceleration was used when available. The entire pipeline was modularized for reproducibility, including data handling, model definition, training logic, and checkpointing. Point cloud encoding and histogram computation were handled using custom PyTorch Dataset and DataLoader classes. Training history, model weights, and learning curves were automatically saved for each run to facilitate experiment tracking and reproducibility.

Hyperparameters are listed in the following table:

| Parameter | Description | Value |
|---|---|---|
| num_points | Number of points randomly sampled per group | 768 |
| bins | Number of bins per axis for histogram | 64 |
| input_shape | Shape of input histogram (C, H, W) | (1, 64, 64) |
| latent_dim | Dimensionality of the latent space | 256 |
| hidden_dims | List of hidden channels in conv layers | [32, 64, 128, 256] |
| lr | Learning rate | 0.03 |
| beta | Weight of the KL divergence in the VAE loss | 0.1 |
| epochs | Number of training epochs | 300 |
| patience | Epochs to wait for early stopping | 5 |
| clip_grad | Max norm for gradient clipping | 1.0 |

**Data visualization**

To qualitatively evaluate the learned latent representation, the output of the bottleneck layer of the trained VAE was visualized. Specifically, the latent representation $\boldsymbol{z}$ was obtained by applying the encoder to the normalized histogram $\boldsymbol{x}$, followed by reparameterization:

$$\boldsymbol{z} = \boldsymbol{\mu} + \boldsymbol{\epsilon} \odot e^{0.5 \cdot \log \boldsymbol{\sigma}^2},$$

where $\boldsymbol{\mu}$ and $\log \boldsymbol{\sigma}^2$ represent the encoder outputs, and $\epsilon \sim \mathcal{N}(\boldsymbol{0}, \boldsymbol{I})$. For visualization purposes, we primarily used $\boldsymbol{\mu}$ as a deterministic representation of the input sample.

The latent vectors were projected onto a 2D space using uniform manifold approximation and projection (UMAP) (version 0.5.5), with n_neighbors=40 to emphasize local neighborhood preservation. This allowed us to assess the clustering tendency and separation of samples in the latent space based on their intrinsic structure.

To further interpret the learned representations, we calculated pairwise cosine similarities between latent vectors and selected sample pairs with either high or low similarity. We then visualized the corresponding input histograms for these pairs, allowing for inspection of how similar or dissimilar representations exhibit in the input space. This approach enabled qualitative validation of whether semantically similar samples were mapped to proximate regions in latent space. Matplotlib 3.7.1 was used for all visualizations.

**Classification of healthy vs cancer samples**

To evaluate the downstream utility of learned representations, a binary classification task was conducted to discriminate between healthy and cancer samples. Latent vectors were extracted by concatenating the mean vectors of two separately trained VAEs (on DPP4 and CD13 histograms). These representations were derived via the encode method from the pretrained models. Data from one center (cohort 1) were used for model training and validation, whereas data from another center (cohort 2) were reserved for testing. To correct for potential batch effects, correlation alignment (CORAL) was applied to match second-order statistics between the training and test features. A variance threshold was used to eliminate near-constant features before classification.

We trained bagging ensembles of support vector machines (SVMs) with different kernels (RBF, polynomial, sigmoid), with hyperparameter optimization performed via Optuna (v3.6.1). The objective was to maximize the cross-validated area under the receiver operating characteristic curve (AUROC) on the training set. The top-k models were then re-trained on the full training set and evaluated on the held-out test set. Final predictions were aggregated using either uniform or performance-weighted ensembling. Model performance was assessed by determining the AUROC and accuracy at the optimal threshold. All experiments were repeated with three different random seeds to ensure robustness.

Hyperparameters are listed in the following table:

| **Parameter** | **Description** | **Value or search space** |
|---|---|---|
| cv | Cross-validation fold | 3 |
| k | Top-k for ensembling | 10 |
| n_trials | Number of trials for hyperparameter optimization | 500 |
| C | Regularization parameter for SVM | [1e-3, 1e2] (log scale) |
| gamma | Kernel coefficient for RBF, poly, and sigmoid kernels | [1e-4, 1.0] (log scale) |
| n_estimators | Number of base SVM classifiers in bagging ensemble | [5, 30] |
| kernel | Kernel type used in SVM | {'rbf', 'poly', 'sigmoid'} |
| degree | Degree of polynomial kernel (if kernel='poly') | [2, 5] |
| coef0 | Independent term in poly/sigmoid kernel | [0.0, 10.0] |

**Supplementary data and comments**

| | | | Healthy controls | Pancreatic tumor |
|---|---|---|---|---|
| Cohort 1 | Category | | Healthy controls | Pancreatic tumor |
| | Sample number | | 31 | 31 |
| | Stage | I | - | 20 |
| | | II | - | 11 |
| | | III | - | 0 |
| | | IV | - | 0 |
| | Age (mean ± S.D.) years old | | 69.7 ± 8.1 | 70.0 ± 9.0 |
| | Gender (female/male) | | 15/16 | 16/15 |
| Cohort 2 | Category | | Healthy controls | Pancreatic tumor |
| | Sample number | | 33 | 29 |
| | Stage | I | - | 4 |
| | | II | - | 25 |
| | | III | - | 0 |
| | | IV | - | 0 |
| | Age (mean ± S.D.) years old | | 36.1 ± 10.8 | 66.7 ± 7.2 |
| | Gender (female/male) | | 16/17 | 8/21 |

**Table S1.** Summary of plasma samples used in this study.

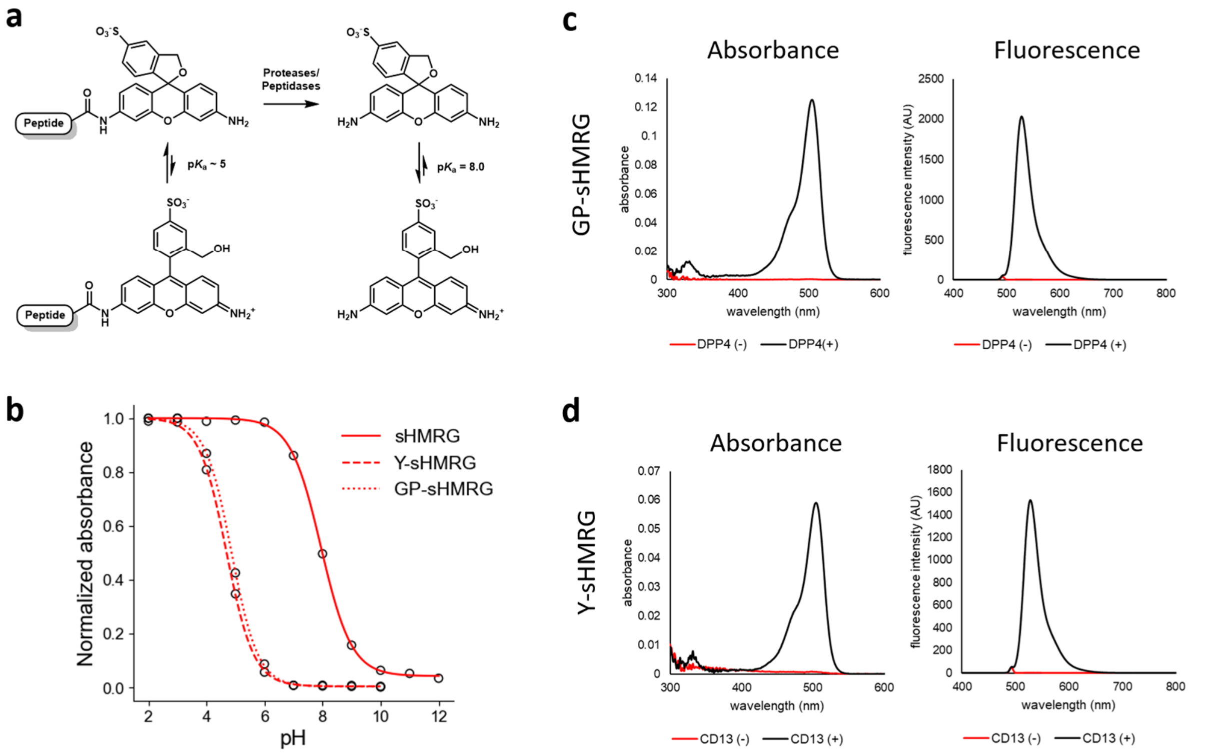


**Figure S1.** Absorbance and fluorescence responses of sHMRG-based fluorogenic probes to target enzymes. (a) Equilibrium of sHMRG and its peptide conjugates. (b) pH-dependent absorbance (500 nm) changes in sHMRG, GP-sHMRG, and Y-sHMRG (3.3 µM) in phosphate buffer (100 mM). (c) Absorbance and fluorescence spectra of GP-sHMRG (3.3 µM) before and after mixing with recombinant DPP4 in HEPES-Na buffer (100 mM, pH 7.4), incubated for 30 min. $\lambda_{ex}$ = 490 nm for fluorescence. (d) Absorbance and fluorescence spectra of Y-sHMRG (3.3 µM) before and after mixing with recombinant CD13 (10 ng/mL) in HEPES-Na buffer (100 mM, pH 7.4), incubated for 30 min. $\lambda_{ex}$ = 490 nm for fluorescence.

**[Comment]**

Absorbance/fluorescence activation of sHMRG-based probes was designed based on the control of the spirolactam-forming equilibrium[5,6]. At neutral pH, amidated sHMRG preferentially assumes the spirolactam form without absorbance in the visible light region. The enzyme reaction cleaving the amide bond generates sHMRG, which preferentially assumes the quinoid form at neutral pH, with an absorbance peak at approximately 500 nm, and becomes fluorescent.

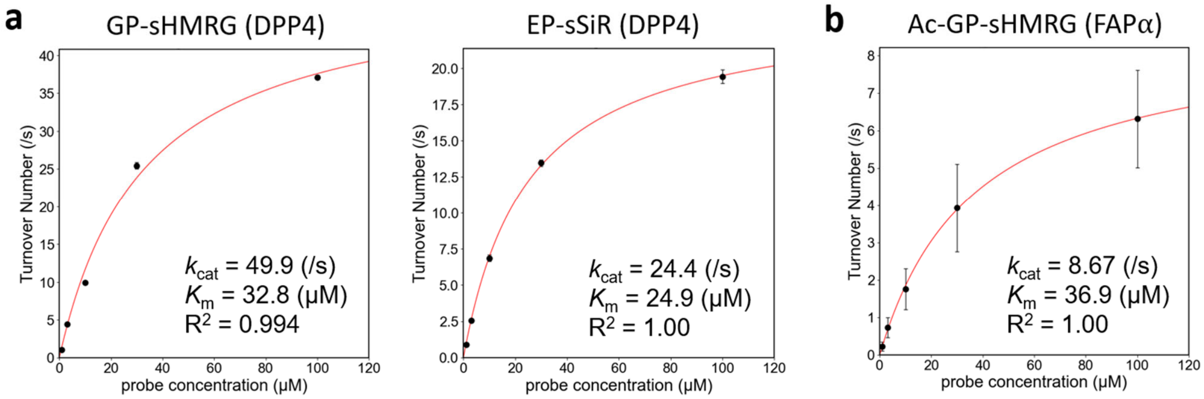


**Figure S2.** M-M plot of GP-sHMRG (left) and EP-sSiR (right) for DPP4 (a) and Ac-GP-sHMRG for FAPα (b). (a) The assay was performed with HEPES-Na buffer (100 mM, pH 7.4) containing $CaCl_2$ (1 mM), $MgCl_2$ (1 mM) and Triton X-100 (150 μM) and DPP4 (25 ng/mL). Fluorescence increase was monitored using multiwell-plate reader, and the initial velocity was converted to concentration change using the fluorescence signal of sHMRG or sSiR (1 μM) as a reference. (b) The assay was performed with HEPES-Na buffer (100 mM, pH 7.4) containing $CaCl_2$ (1 mM), $MgCl_2$ (1 mM) and Triton X-100 (150 μM) and FAPα (10 ng/mL).

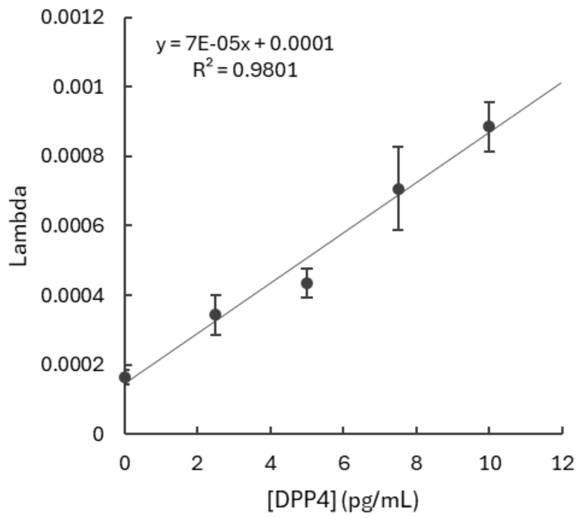


**Figure S3.** Detection of single-molecule activity of DPP4 using the COP-based microdevice. GP-sHMRG (10 μM), DPP4 (0-10 pg/mL) in HEPES-Na buffer (100 mM, pH 7.4) containing $CaCl_2$ (1 mM), $MgCl_2$ (1 mM), DTT (100 μM), and Triton X-100 (150 μM) was loaded into the COP-based microdevice, sealed, and incubated for 2 h at 25°C. Lambda (probability of an enzyme molecule being present in a well) values were calculated by dividing the number of activity spots by the total number of wells analyzed. Error bars represent SD (n = 3).

**[Comment]**

As proof of that bright spots reflect the number of molecules in a sample, we confirmed that the number of bright spots reflects the concentration of DPP4, whereas no significant change in the fluorescence intensity of bright spots was observed[7,8]. Limit of detection (LOD) was calculated to be 1.18 pg/mL (3σ), and limit of quantification (LOQ) was calculated to be 3.15 pg/mL (10σ).

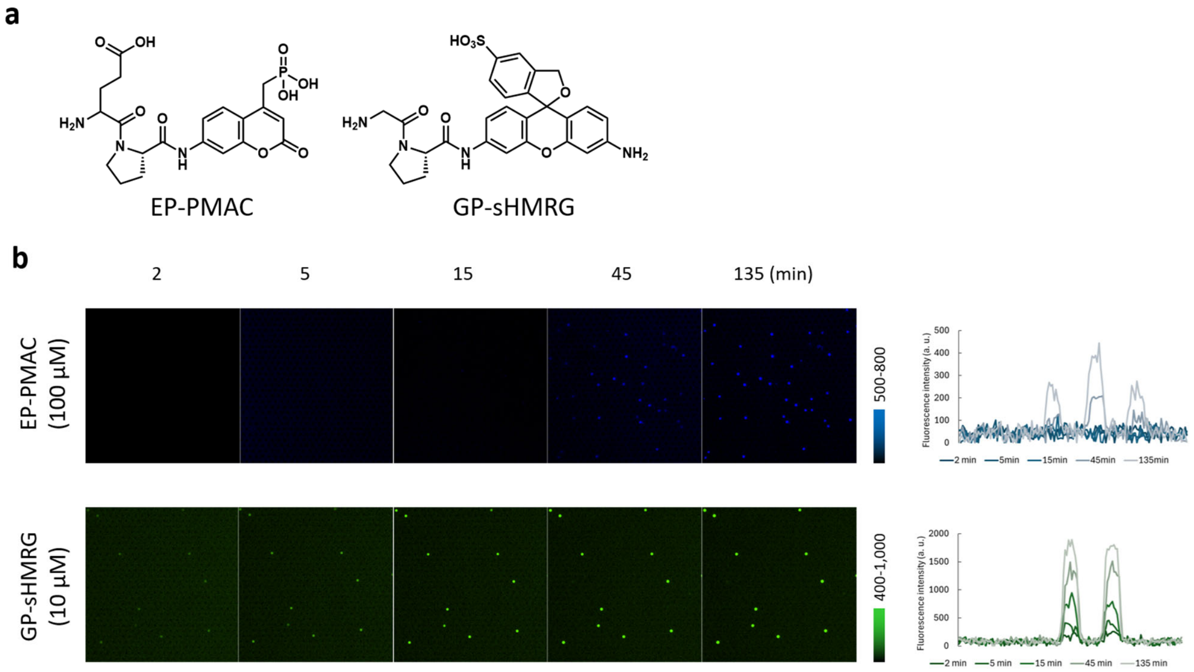


**Figure S4.** Detection of single-molecule DPP4 activity using FRAD. (a) Molecular structures of EP-PMAC and GP-sHMRG. (b) Fluorescence images of the microdevice after loading GP-sHMRG (10 μM) or EP-PMAC (100 μM), DPP4 (0.1 ng/mL) in HEPES-Na buffer (100 mM, pH 7.4) containing $CaCl_2$ (1 mM), $MgCl_2$ (1 mM), DTT (100 μM), and Triton X-100 (3 mM), incubated for indicated times at 25°C. Representative intensity profiles of the bright spots are shown to the right.

**[Comment]** Detection of single-molecule DPP4 activity was compared between sHMRG-based probes and PMAC-based probes[9]. The same microdevice with a glass coverslip bottom (FRAD) was used. In sHMRG-based probe, bright spots were detectable within 5 min using 10 μM probe, whereas the PMAC-based probe provided a detectable signal only after 45 min even when 100 μM probe was used. In sHMRG-based probe, the signal reached a plateau at 135 min, reflecting consumption of the substrate, so in the following experiments, probe concentration was set to 30 μM to minimize the effect.

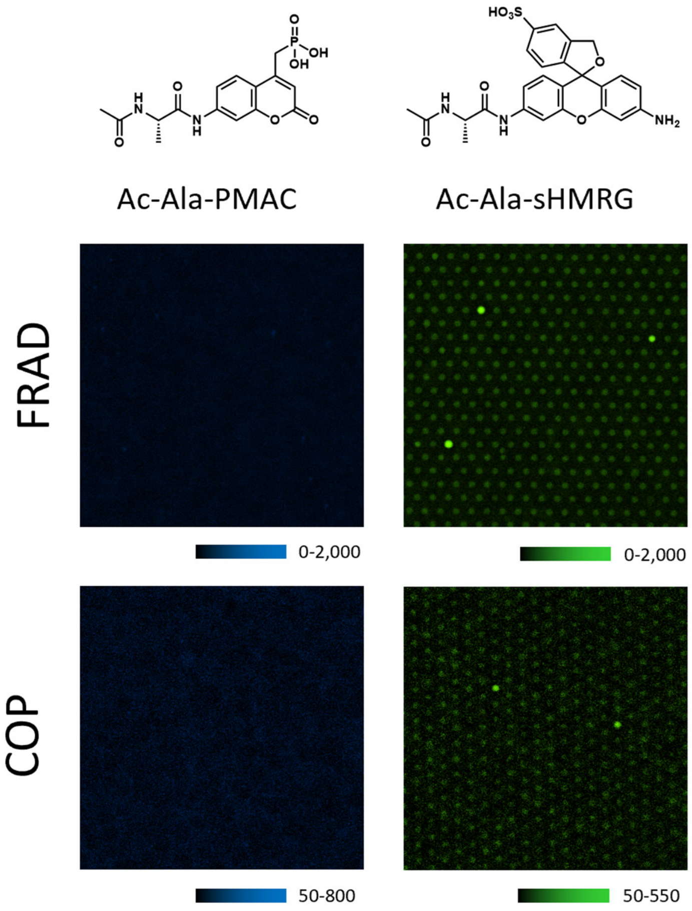


**Figure S5.** Detection of single-molecule APEH activity in blood using FRAD and the COP microdevice. Figure shows fluorescence images of the microdevice (FRAD or COP) after loading Ac-Ala-sHMRG (10 μM) or Ac-Ala-PMAC (50 μM), blood samples of healthy human subjects (1/1000) in HEPES-Na buffer (100 mM, pH 7.4) containing $CaCl_2$ (1 mM), $MgCl_2$ (1 mM), DTT (100 μM), and Triton X-100 (3 mM or FRAD and 250 μM for COP microdevice), incubated for 2 h at 25°C.

**[Comment]** In an attempt to detect the activity of acylamino acid–releasing enzyme (APEH) activity in blood samples, Ac-Ala, a substrate sequence of APEH[10], was attached to sHMRG and PMAC, and the signals were compared for FRAD- and COP microdevice–based single-molecule analysis. Only the sHMRG-based probe was able to detect the desired activity. We consider that this reflected the weak penetration of light and weak brightness of PMAC.

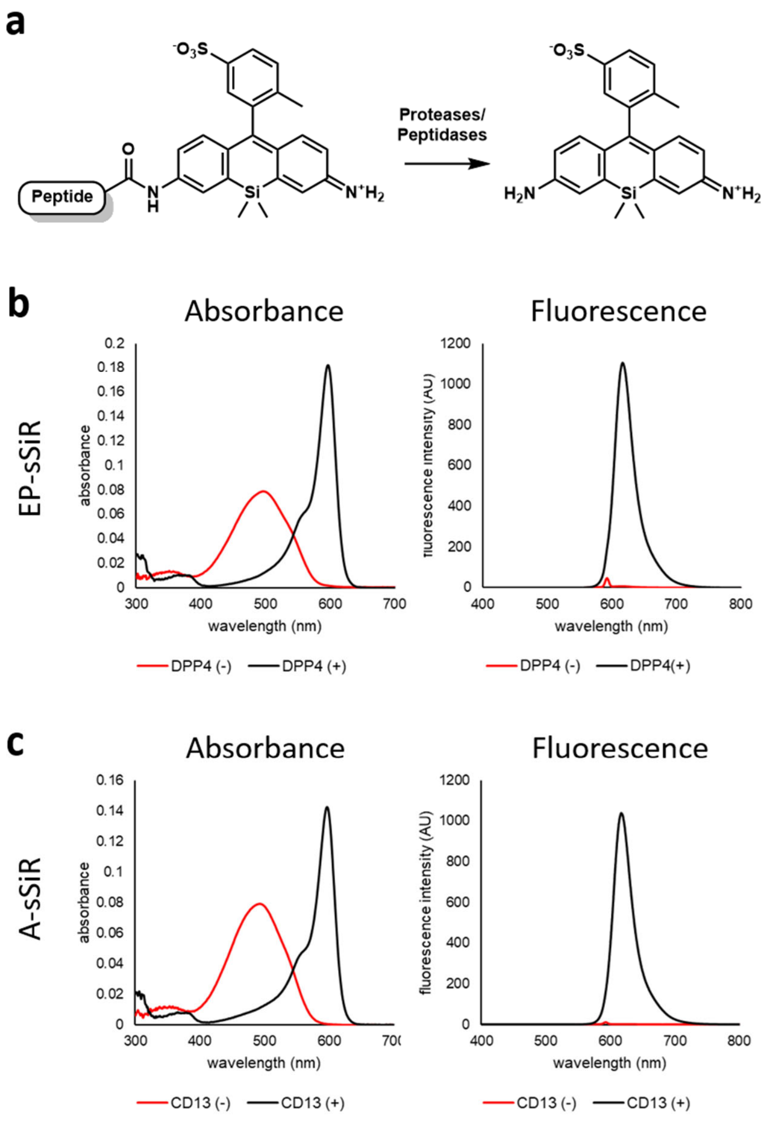


**Figure S6.** Absorbance and fluorescence responses of sSiR-based fluorogenic probes to target enzymes. (a) Structures of sHMRG and peptide conjugates. (b) Absorbance and fluorescence spectra of EP-SiR (3.3 μM) before and after mixing with recombinant DPP4 in HEPES-Na buffer (100 mM, pH 7.4) and incubating for 90 min. $\lambda_{ex}$ = 590 nm for fluorescence. (c) Absorbance and fluorescence spectra of A-sSiR (3.3 μM) before and after mixing with recombinant CD13 (10 ng/mL) in HEPES-Na buffer (100 mM, pH 7.4) and incubating for 90 min. $\lambda_{ex}$ = 590 nm for fluorescence.

**[Comment]** Silicon-rhodamine (SiR) exhibits an absorbance/fluorescence wavelength shift upon amidation of one of the anilines of rhodamine[11,12], and we exploited this characteristic to design fluorogenic probes for proteases/peptidases. The introduction of sulfonic acid to increase retention in the microdevice did not alter this characteristic, and upon excitation at 590 nm to selectively excite non-amidated sSiR, fluorescence activation was clearly observed during peptide-cleaving reactions.

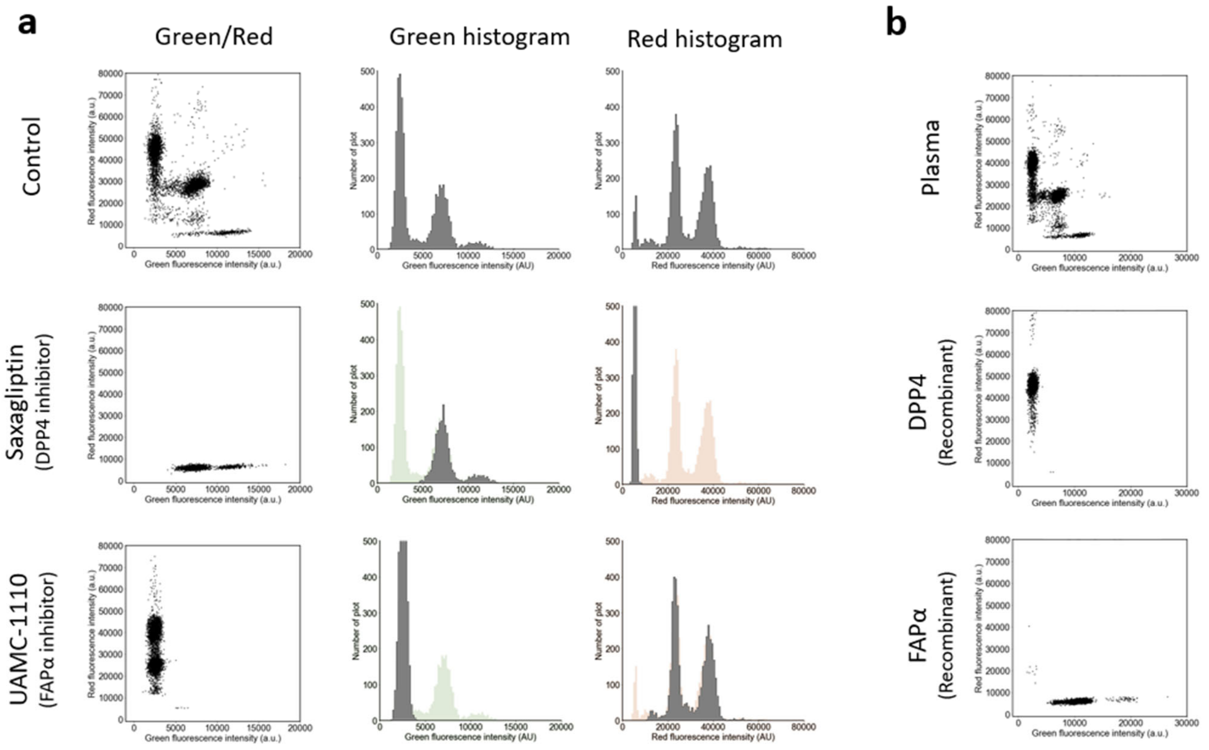


**Figure S7.** Identification of DPP4 and FAPα activity. (a) Dot plots and histograms of the fluorescence intensity of microwells after loading Ac-GP-sHMRG (green, FAPα substrate, horizontal axis, 30 μM), EP-sSiR (red, DPP4 substrate, vertical axis, 30 μM), IRDye800 (30 μM), and blood samples from healthy human subjects (1/5,000 dilution) in HEPES-Na buffer (100 mM, pH 7.4), $CaCl_2$ (1 mM), $MgCl_2$ (1 mM), DTT (100 μM), Triton X-100 (150 μM), and BSA (1% w/v), incubated at 25°C for 4 h. For inhibitors, saxagliptin (DPP4 inhibitor, 100 nM) or UAMC-1110 (FAPα inhibitor, 32 nM) were used. The histogram of the control condition was overlayed (as pale green or orange) on histograms for inhibitor treatments for comparison. (b) Dot plots of the fluorescence intensity of microwells after loading Ac-GP-sHMRG (30 μM), EP-sSiR (30 μM), IRDye800 (10 μM), and recombinant DPP4 or FAPα (0.2 ng/mL) in HEPES-Na buffer (100 mM, pH 7.4), $CaCl_2$ (1 mM), $MgCl_2$ (1 mM), DTT (100 μM), Triton X-100 (150 μM), and BSA (1% w/v), incubated for 4 h.

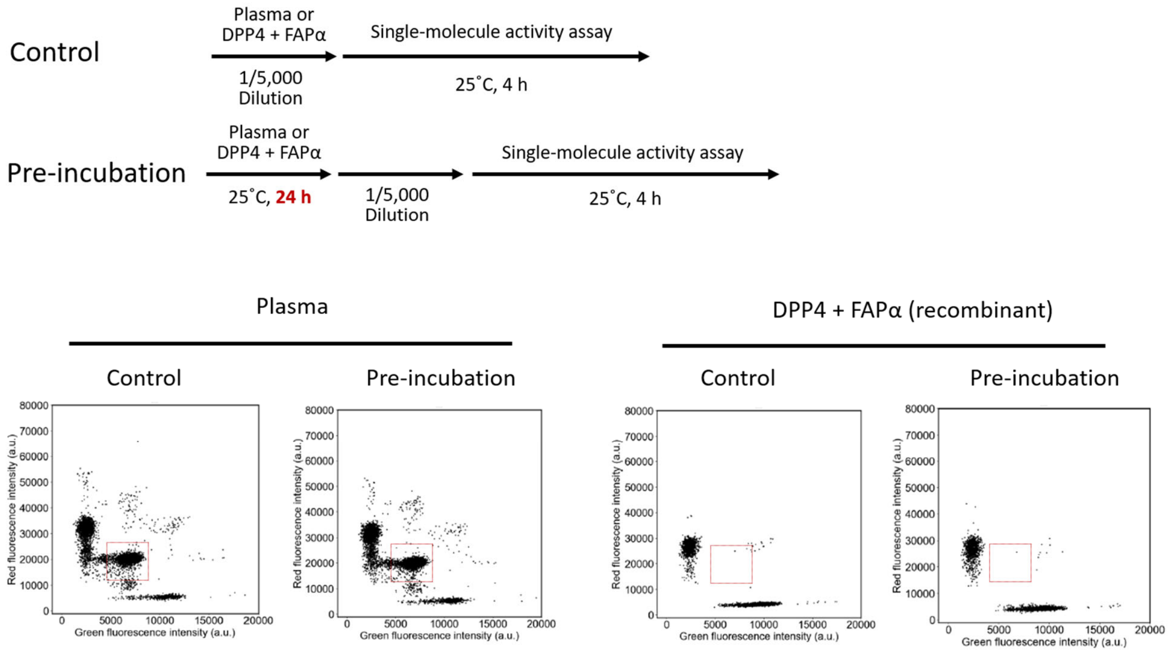


**Figure S8.** Equilibrium between DPP4 and FAPα. Plasma samples or a mixture of recombinant DPP4 and FAPα (0.1 ng/mL each) were incubated in HEPES-Na buffer (100 mM, pH 7.4), $CaCl_2$ (1 mM), $MgCl_2$ (1 mM), DTT (100 μM), Triton X-100 (150 μM), and BSA (1% w/v) at 25°C for 24 h. The samples were then diluted 1/5,000 in the same buffer, and the activity was analyzed using the same conditions used for experiments shown in **Figure S7**.

**[Comment]** In order to evaluate the equilibrium of DPP4-FAPα heterodimer formation, we mixed recombinant DPP4 (homodimer) and FAPα (homodimer) at high concentrations and incubated the mixtures for 24 h to test if the heterodimer would be generated. No cluster corresponding to the DPP4-FAPα heterodimer was observed, indicating that homodimers of DPP4 and FAPα were not in equilibrium with the DPP4-FAPα heterodimer, at least under the condition and timeframe tested.

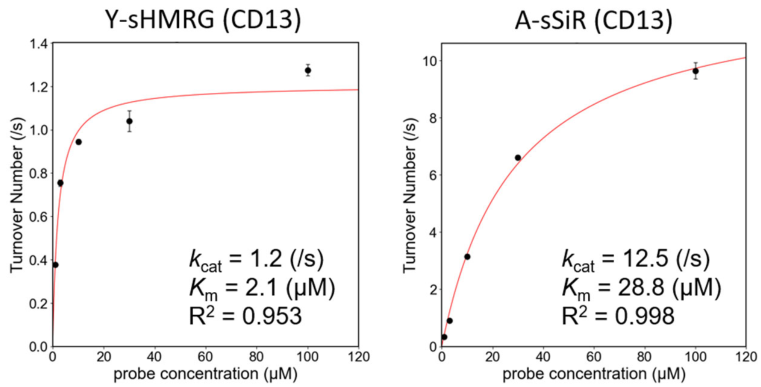


**Figure S9.** M-M plot of Y-sHMRG (left) and A-sSiR (right) for DPP4. The assay was performed with HEPES-Na buffer (100 mM, pH 7.4) containing $CaCl_2$ (1 mM), $MgCl_2$ (1 mM) and Triton X-100 (150 μM) and DPP4 (25 ng/mL). Fluorescence increase was monitored using multiwell-plate reader, and the initial velocity was converted to concentration change using the fluorescence signal of sHMRG or sSiR (1 μM) as a reference.

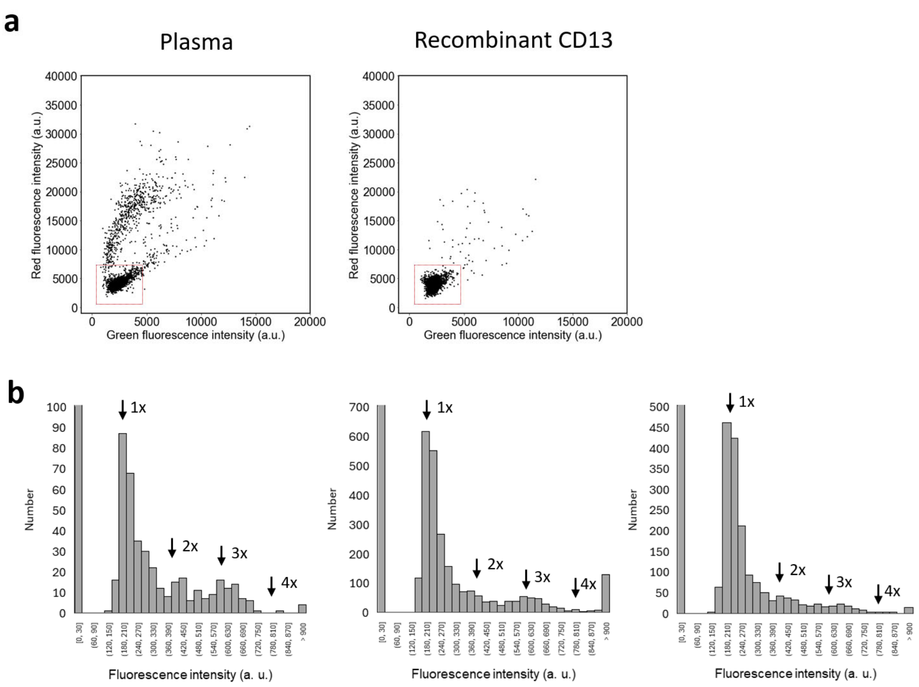


**Figure S10.** Identification of CD13 activity. (a) Dot plots of the fluorescence intensity of microwells after loading Y-sHMRG (green, horizontal axis, 30 μM), A-sSiR (red, vertical axis, 30 μM), IRDye800 (10 μM), and blood samples from healthy human subjects (1/5,000 dilution, left) or recombinant CD13 (0.2 ng/mL, right) in HEPES-Na buffer (100 mM, pH 7.4), $CaCl_2$ (1 mM), $MgCl_2$ (1 mM), DTT (100 μM) and Triton X-100 (150 μM), incubated at 25°C for 2 h. (b) Histograms of CD13 activity in three representative PDAC patients. Fluorescence intensity of the FITC channel is shown, and 1× indicates the fluorescence intensity corresponding to that of recombinant human CD13, whereas 2-4× shows the intensity multiple of 1×.

**[Comment]** In samples from some PDAC patients, bright spots exhibiting high activity (corresponding to cluster [3] of **Figure 2d**) were observed. As the data did not follow Poisson's distribution, this result seems to reflect the presence of molecular species consisting of multiple CD13 molecules. Major active species exhibited 2-3 times the activity of recombinant CD13, but species with higher activity than 4× were also detected, indicating that oligomer species were present as well.

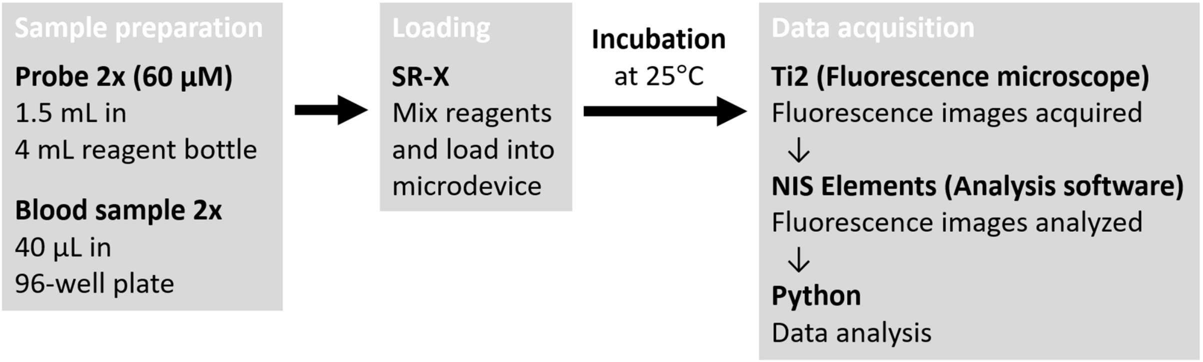


**Figure S11.** Flow of analysis for automated data acquisition.

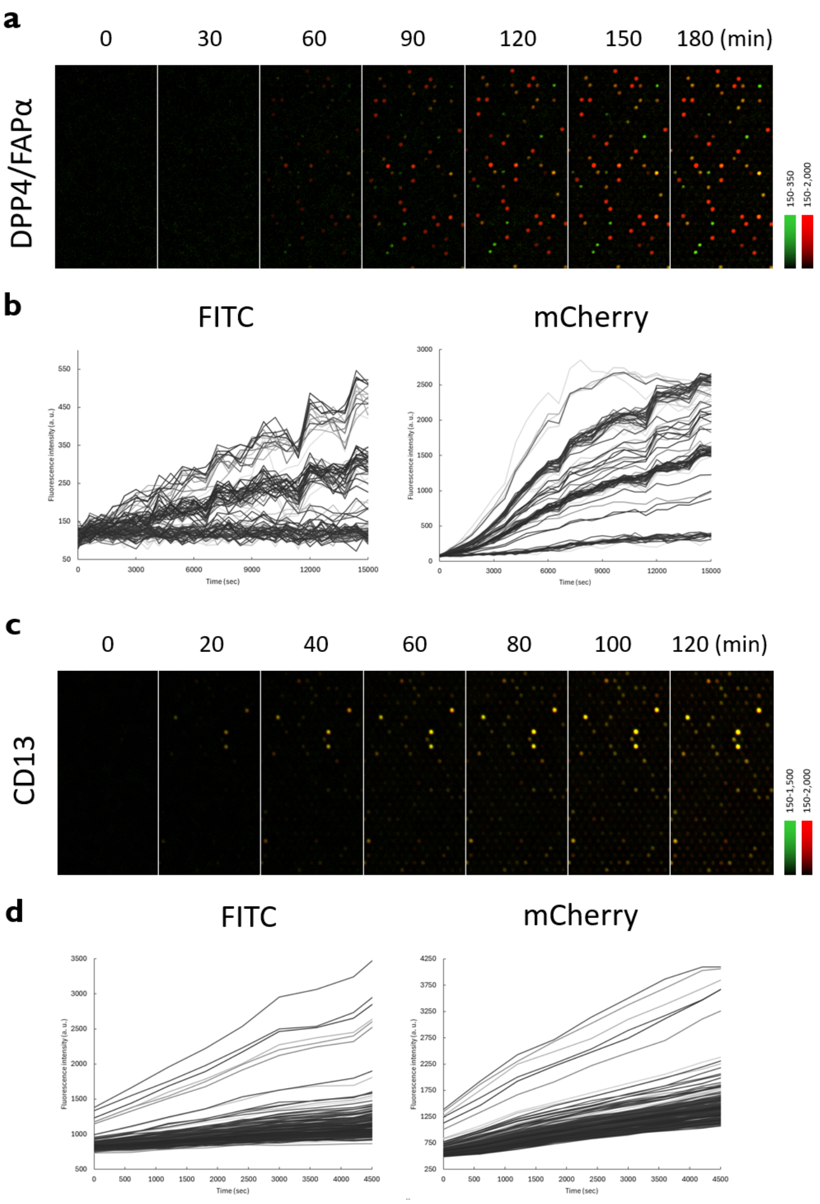


**Figure S12.** Time course of fluorescence increase of wells in the optimized assay conditions. (a) Fluorescence images of microdevice after loading Ac-GP-sHMRG (30 μM) and EP-sSiR (30 μM), blood samples of healthy human subjects (1/1000) in HEPES-Na buffer (100 mM, pH 7.4) containing $CaCl_2$ (1 mM), $MgCl_2$ (1 mM), DTT (100 μM), Triton X-100 (150 μM) and BSA (1% w/v) and incubated at 25°C for indicated time. (b) Quantification of fluorescent intensities at green (FITC) and red (mCherry) channels of 100 activity spots in (a). (c) Fluorescence images of microdevice after loading Y-sHMRG (30 μM) and A-sSiR (30 μM), blood samples of PDAC patients with high CD13 count (1/2000) in HEPES-Na buffer (100 mM, pH 7.4) containing $CaCl_2$ (1 mM), $MgCl_2$ (1 mM), DTT (100 μM) and Triton X-100 (150 μM) and incubated at 25°C for indicated time. (d) Quantification of fluorescent intensities at green (FITC) and red (mCherry) channels of 250 activity spots in (c).

**a**

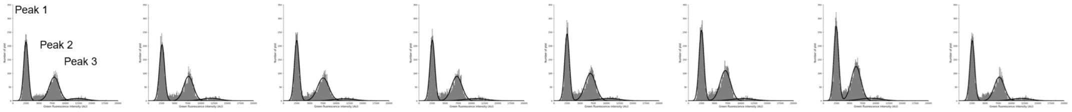


| Data | Peak 1 F. I. | Peak 1 FWHM | Peak 1 Lambda | Peak 2 F. I. | Peak 2 FWHM | Peak 2 Lambda | Peak 3 F. I. | Peak 3 FWHM | Peak 3 Lambda | High activity population | Total Lambda | F. I. CV between 12 images (%) | Lambda CV between 12 images (%) |
|---|---|---|---|---|---|---|---|---|---|---|---|---|---|
| #1 | 2512 | 1047 | 0.0280 | 7881 | 2227 | 0.0261 | 12331 | 4104 | 0.0052 | 0.52 | 0.0593 | 2.3 | 2.8 |
| #2 | 2597 | 1142 | 0.0292 | 7643 | 2174 | 0.0258 | 11879 | 4040 | 0.0055 | 0.53 | 0.0606 | 3.8 | 4.2 |
| #3 | 2528 | 1080 | 0.0295 | 7530 | 2524 | 0.0272 | 11520 | 4659 | 0.0049 | 0.52 | 0.0615 | 3.0 | 5.3 |
| #4 | 2563 | 1106 | 0.0313 | 7212 | 2347 | 0.0280 | 10707 | 4999 | 0.0054 | 0.53 | 0.0647 | 2.8 | 5.6 |
| #5 | 2505 | 1073 | 0.0325 | 6940 | 2382 | 0.0303 | 10370 | 4802 | 0.0054 | 0.52 | 0.0682 | 2.9 | 5.8 |
| #6 | 2510 | 1100 | 0.0348 | 7041 | 2321 | 0.0330 | 10403 | 5189 | 0.0053 | 0.51 | 0.0732 | 2.0 | 5.0 |
| #7 | 2466 | 1087 | 0.0365 | 6240 | 2181 | 0.0337 | 9053 | 4873 | 0.0056 | 0.52 | 0.0758 | 2.4 | 2.5 |
| #8 | 2556 | 1060 | 0.0289 | 7691 | 2255 | 0.0264 | 11904 | 4279 | 0.0054 | 0.52 | 0.0607 | 1.3 | 5.6 |
| CV (%) | 1.6 | 2.7 | 9.7 | 7.3 | 5.1 | 10.9 | 9.9 | 9.3 | 4.1 | 1.1 | 9.6 | | |

**b**

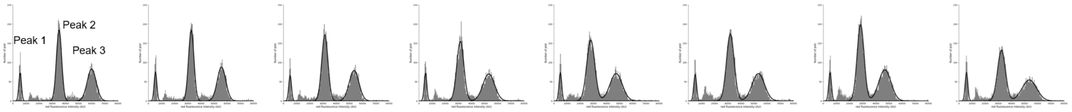


| Data | Peak 1 F. I. | Peak 1 FWHM | Peak 1 Lambda | Peak 2 F. I. | Peak 2 FWHM | Peak 2 Lambda | Peak 3 F. I. | Peak 3 FWHM | Peak 3 Lambda | High activity population | Total Lambda | F. I. CV between 12 images (%) | Lambda CV between 12 images (%) |
|---|---|---|---|---|---|---|---|---|---|---|---|---|---|
| #1 | 5591 | 2343 | 0.0052 | 35185 | 4714 | 0.0314 | 59998 | 7999 | 0.0227 | 0.42 | 0.0593 | 1.8 | 2.8 |
| #2 | 5429 | 2396 | 0.0055 | 32617 | 4845 | 0.0322 | 55659 | 7665 | 0.0229 | 0.41 | 0.0606 | 2.0 | 4.2 |
| #3 | 5068 | 2426 | 0.0049 | 31372 | 5585 | 0.0343 | 53529 | 8614 | 0.0224 | 0.39 | 0.0615 | 2.3 | 5.3 |
| #4 | 5013 | 2412 | 0.0054 | 31515 | 6342 | 0.0353 | 53116 | 10450 | 0.0240 | 0.40 | 0.0647 | 2.9 | 5.6 |
| #5 | 4940 | 2409 | 0.0054 | 28057 | 7232 | 0.0384 | 47321 | 11074 | 0.0244 | 0.39 | 0.0682 | 7.2 | 5.8 |
| #6 | 5255 | 2506 | 0.0053 | 31982 | 6846 | 0.0426 | 53295 | 11100 | 0.0252 | 0.37 | 0.0732 | 3.5 | 5.0 |
| #7 | 5202 | 2463 | 0.0056 | 28397 | 6252 | 0.0429 | 46847 | 10891 | 0.0273 | 0.39 | 0.0758 | 3.8 | 2.5 |
| #8 | 5819 | 2394 | 0.0054 | 32438 | 7064 | 0.0334 | 53638 | 13040 | 0.0218 | 0.40 | 0.0607 | 5.6 | 5.6 |
| CV (%) | 5.7 | 2.0 | 4.1 | 7.4 | 15.9 | 12.4 | 8.0 | 18.3 | 7.6 | 3.9 | 9.6 | | |

**Figure S13.** Repeatability of the single-molecule enzyme assay to detect DPP4/FAPα activity. Analysis of blood samples were performed eight times in the same condition as **Figure 2a**. (a) The histograms of Ac-GP-sHMRG (30 μM) were fitted to Gaussian distributions. Fluorescence intensities of the peak top, full width at half maximum (FWHM), and number of spots were calculated for each peak. (b) The histograms of EP-sSiR were fitted to Gaussian distributions.

**a**

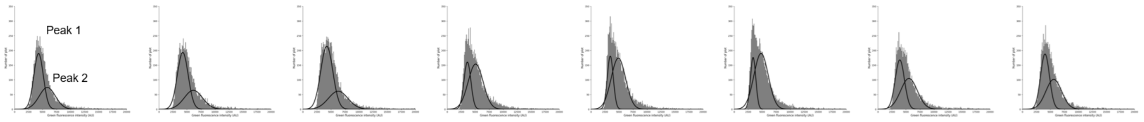


| Data | Peak 1 F. I. | Peak 2 F. I. | Total Lambda | F. I. CV between 9 images (%) | Lambda CV between 9 images (%) |
|---|---|---|---|---|---|
| #1 | 4287 | 5848 | 0.0853 | 3.8 | 5.5 |
| #2 | 4244 | 6145 | 0.0852 | 2.9 | 7.8 |
| #3 | 4252 | 6281 | 0.0978 | 4.4 | 7.4 |
| #4 | 3606 | 5081 | 0.0943 | 2.1 | 8.2 |
| #5 | 3447 | 4888 | 0.0982 | 4.1 | 6.1 |
| #6 | 3388 | 4810 | 0.0999 | 1.6 | 4.4 |
| #7 | 3902 | 5539 | 0.0877 | 1.8 | 4.8 |
| #8 | 4039 | 5601 | 0.0867 | 3.1 | 4.3 |
| CV (%) | 9.5 | 10.1 | 6.9 | | |

**b**

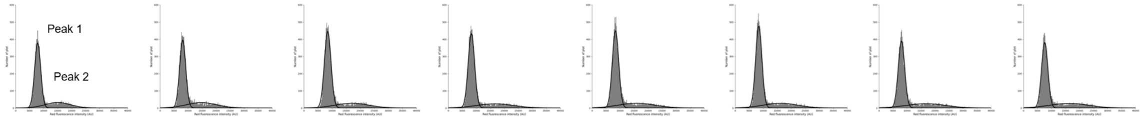


| Data | Peak 1 F. I. | Peak 2 F. I. | Total Lambda | F. I. CV between 9 images (%) | Lambda CV between 9 images (%) |
|---|---|---|---|---|---|
| #1 | 7874 | 15041 | 0.0853 | 6.0 | 5.5 |
| #2 | 8086 | 14789 | 0.0852 | 3.8 | 7.8 |
| #3 | 8462 | 17099 | 0.0978 | 2.9 | 7.4 |
| #4 | 8265 | 16738 | 0.0943 | 2.7 | 8.2 |
| #5 | 8377 | 15945 | 0.0982 | 2.1 | 6.1 |
| #6 | 8416 | 16153 | 0.0999 | 2.3 | 4.4 |
| #7 | 8021 | 16743 | 0.0877 | 3.3 | 4.8 |
| #8 | 7599 | 16315 | 0.0867 | 3.4 | 4.3 |
| CV (%) | 3.7 | 5.1 | 6.9 | | |

**Figure S14.** Repeatability of the single-molecule enzyme assay to detect CD13 activity. Analysis of blood samples were performed eight times in the same condition as **Figure 2c**. (a) The histograms of Y-sHMRG (30 μM) were fitted to Gaussian distributions. Fluorescence intensities of the peak top, full width at half maximum (FWHM), and number of spots were calculated for each peak. (b) The histograms of A-sSiR were fitted to Gaussian distributions.

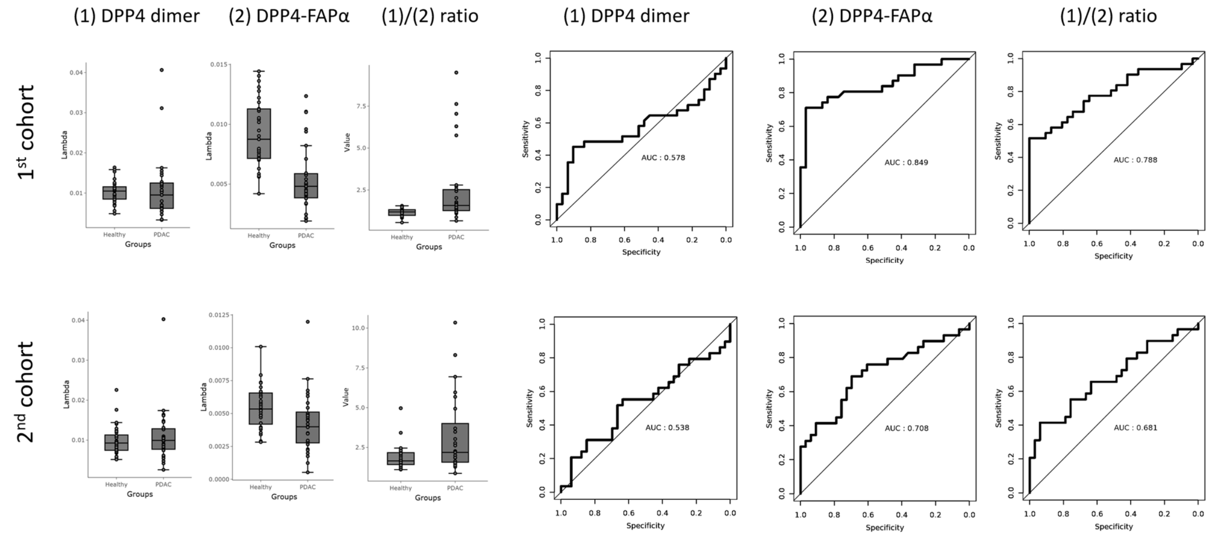


**Figure S15.** Clusters of DPP4 for PDAC diagnosis. Dot plots and ROC curves are shown for 1st (top) and 2nd (bottom) cohort. The thresholding follows that of **Figure 2b**.

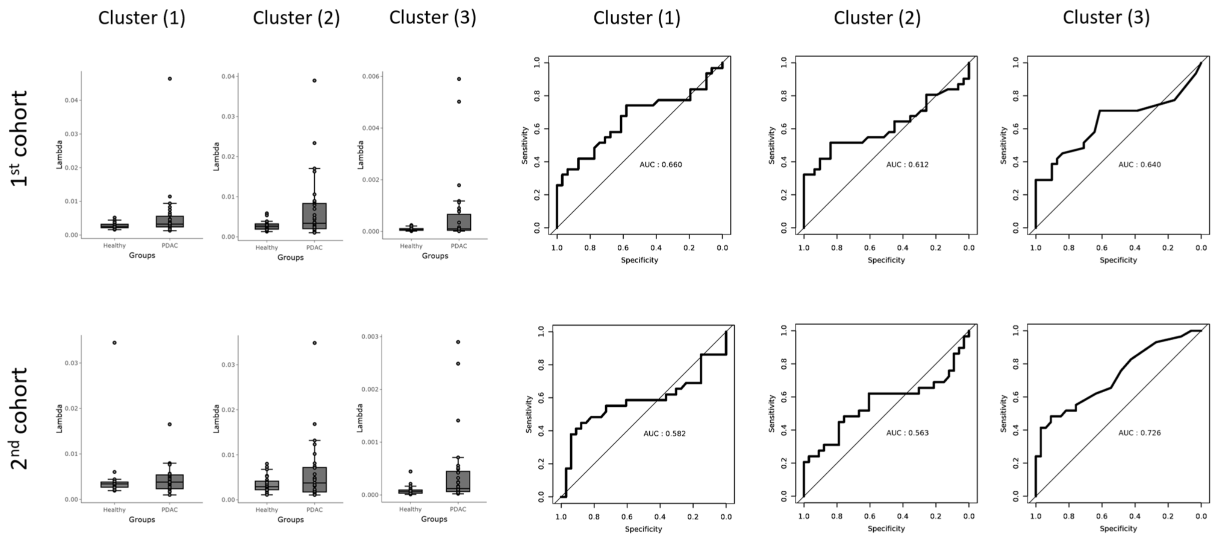


**Figure S16.** Clusters of CD13 for PDAC diagnosis. Dot plots and ROC curves are shown for 1st (top) and 2nd (bottom) cohort. The thresholding follows that of **Figure 2d**.

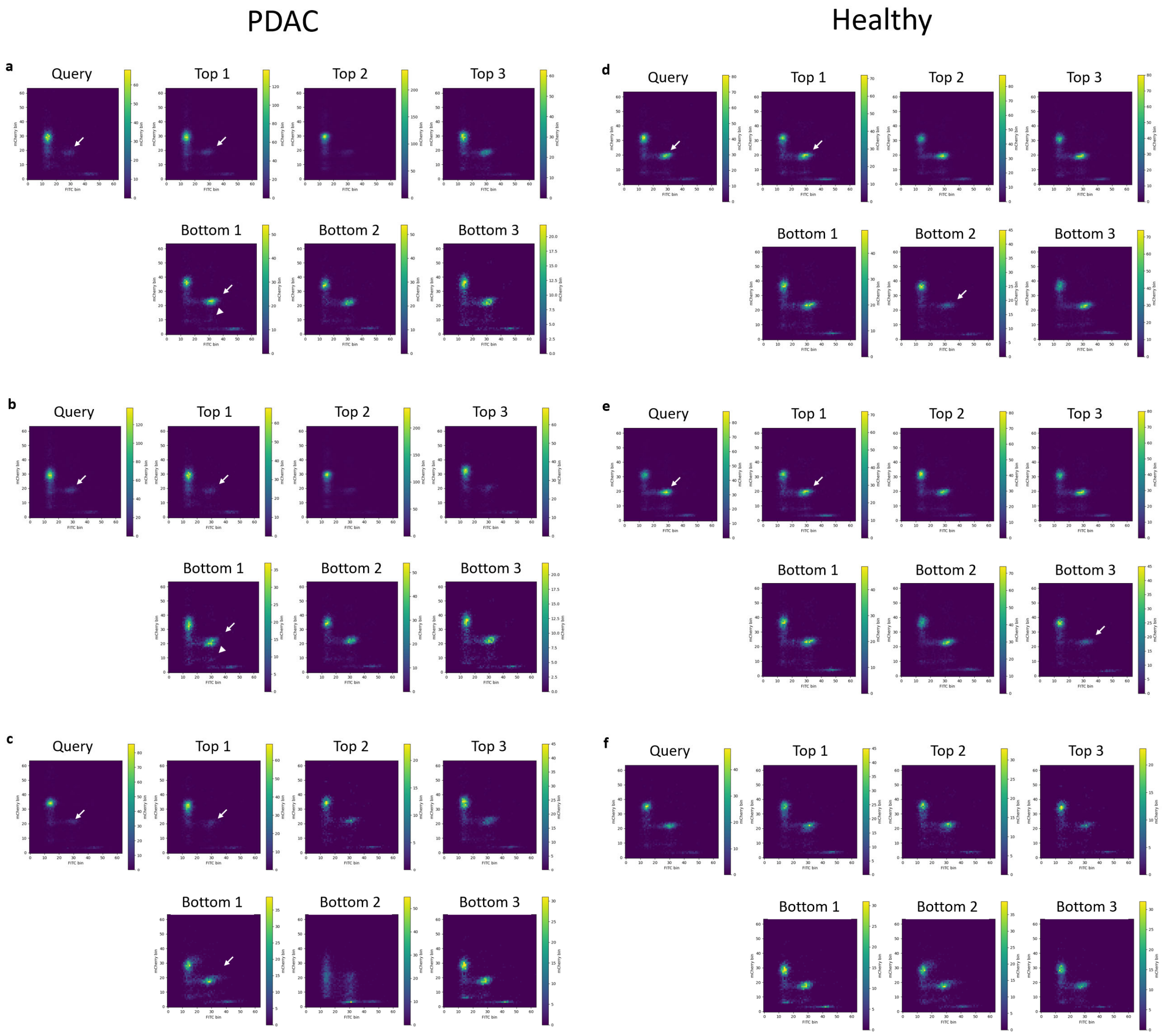


**Figure S17.** Representative examples of top 3 and bottom 3 likely specimen for each query sample for the analysis of DPP4. (a)-(c) are representative data of PDAC patients, and (d)-(f) are representative data of healthy subjects.

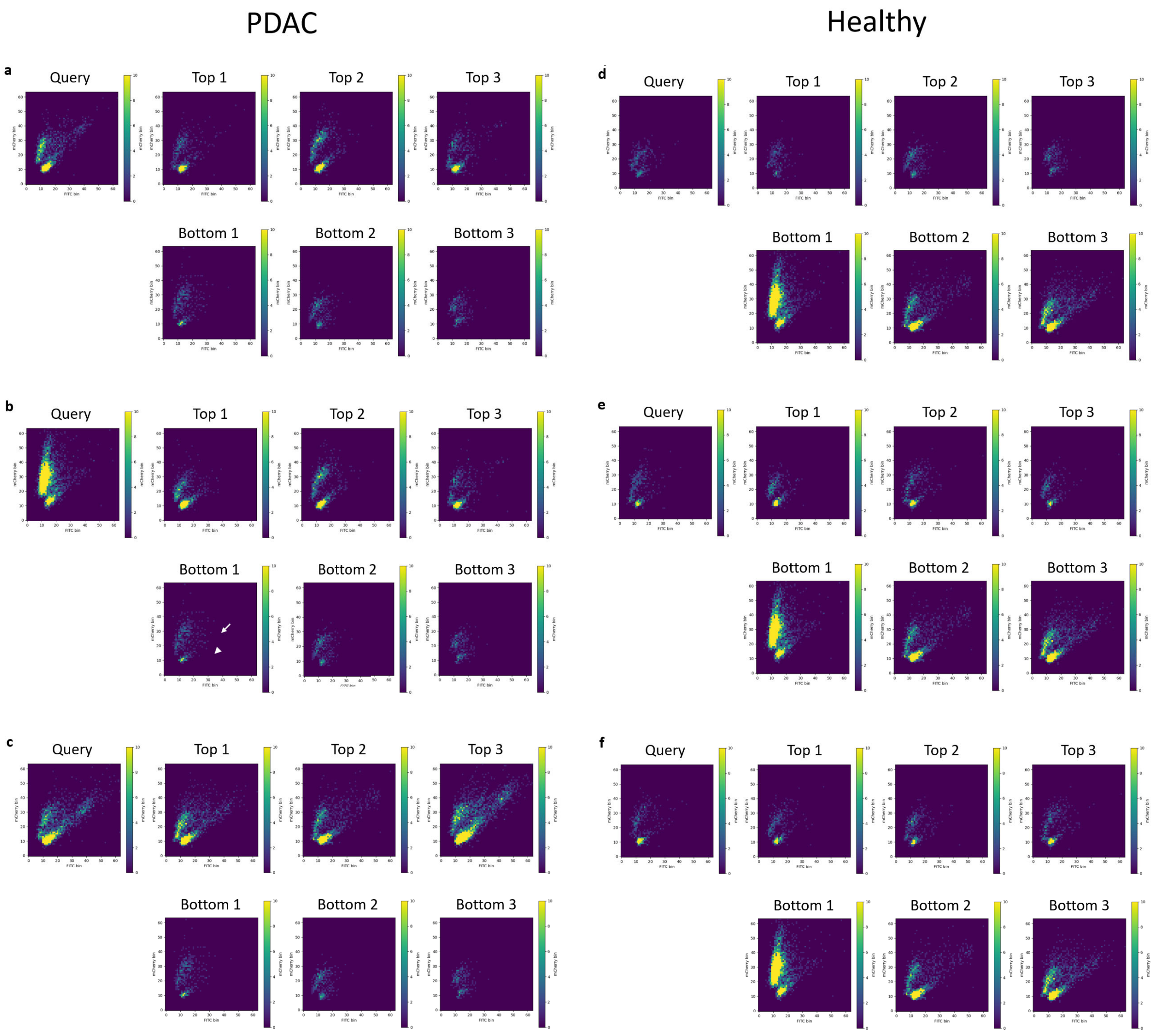


**Figure S18**. Representative examples of top 3 and bottom 3 likely specimen for each query sample for the analysis of CD13. (a)-(c) are representative data of PDAC patients, and (d)-(f) are representative data of healthy subjects.

### Supplementary data for synthesis and characterization of compounds

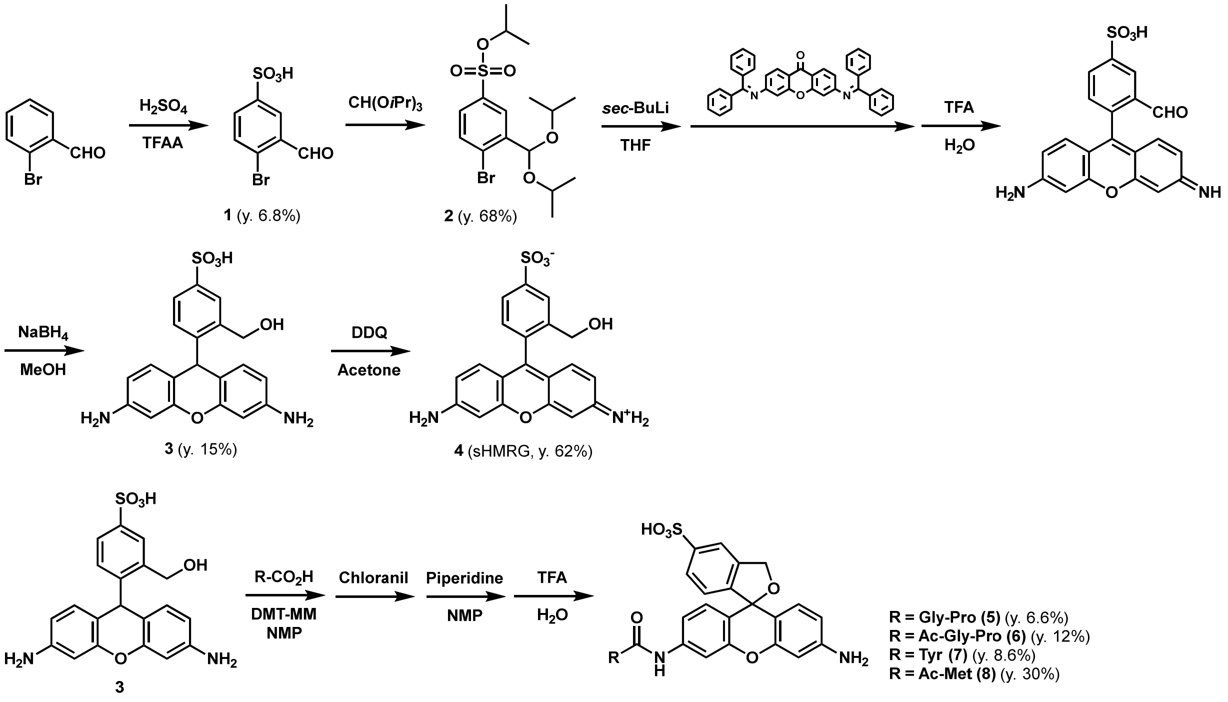


**Scheme S1.** Preparation of sHMRG and fluorogenic probes for proteases/peptidases.

#### Preparation of 4-bromo-3-formylbenzenesulfonic acid (1)

Trifluoroacetic anhydride (TFAA; 15.7 mL, 113.6 mmol) was added to $H_2SO_4$ (2.88 mL, 54.1 mmol) with cooling at 0°C under Ar, then the solution was brought to room temperature and stirred for 3 hours. 2-Bromobenzaldehyde (6.25 mL, 54.1 mmol) was added dropwise under cooling at 0°C, then the solution was brought to room temperature and stirred for 17 h. Water was added dropwise under cooling at 0°C, and TFAA and water were removed in vacuo. The residue was purified by MPLC (eluent: A/B = 95/5 to 5/95, A: 100% $H_2O$ containing 0.1% trifluoroacetic acid (TFA), B: 100% $CH_3CN$ containing 0.1% TFA) and eluent was freeze-dried to afford **1** (982 mg, 3.71 mmol) as a colorless solid, yield 6.8%.

$^1$H-NMR (400 MHz, DMSO-$d_6$) δ 10.17 (s, 1H), 7.98 (m, 1H), 7.74 (m, 2H).

$^{13}$C-NMR (100 MHz, DMSO-$d_6$) δ 192.0, 148.9, 134.3, 133.2, 133.1, 127.4, 126.1.

HRMS (ESI$^-$): *m/z* Calcd. for [M-H]$^-$, 262.90137, Found, 262.89719 (-4.17 mDa)

#### Preparation of isopropyl 4-bromo-3-(diisopropoxymethyl)benzenesulfonate (2)

A solution of 4-bromo-3-formylbenzenesulfonic acid (338 mg, 1.28 mmol) in triisopropyl orthoformate (2.8 mL, 12.8 mmol) was stirred at 60°C for 1 h, and then evaporated. The residue was purified by column chromatography on silica gel (eluent: AcOEt/hexane = 0/100 > 20/80) to afford **2** (357 mg, 0.87 mmol) as a colorless solid, yield 68%.

$^1$H-NMR (400 MHz, $CDCl_3$) δ 8.22 (m, 1H), 7.68 (m, 2H), 4.76 (sep, *J* = 5.9 Hz, 1H), 3.89 (sep, *J* = 6.2 Hz, 2H), 1.28 (d, *J* = 5.9 Hz, 6H), 1.18 (dd, *J* = 13.0 Hz, 6.2 Hz, 12H)

$^{13}$C-NMR (100 MHz, $CDCl_3$) δ 141.2, 137.0, 133.7, 128.7, 128.5, 128.1, 97.9, 69.3, 23.0, 22.9, 22.5.
HRMS (ESI$^+$): *m/z* Calcd. for [M+Na]$^+$, 431.05038, Found, 431.04944 (-0.94 mDa).

**Preparation of leuco-sHMRG (3)**

A solution of isopropyl 4-bromo-3-(diisopropoxymethyl)benzenesulfonate (213 mg, 0.52 mmol) in THF was cooled to -78°C, and then 1.3 M *sec*-BuLi (0.4 mL, 0.52 mmol) was added dropwise. The mixture was stirred for 30 min, then a solution of 3,6-bis-(diphenylmethyleneamino)xanthone (45 mg, 0.081 mmol) in THF was slowly added, and the whole was brought to room temperature and further stirred for 1.5 h. Next, TFA and water were added and the reaction mixture was stirred for 3.5 h, and then evaporated. The residue was purified by MPLC (eluent: A/B = 95/5 to 5/95, A: 100% $H_2O$ containing 0.1% TFA, B: 100% $CH_3CN$ containing 0.1% TFA) and eluent was freeze-dried to afford the formyl product as a crude.

The crude (130 mg) was dissolved in MeOH and $NaBH_4$ (90 mg, 2.38 mmol) was added slowly under cooling at 0°C and stirred for 30 min at 0°C, and then evaporated. The residue was purified by MPLC (eluent: A/B = 95/5 to 5/95, A: 100% $H_2O$ containing 0.1M triethylamine acetate (TEAA), B: 100% $CH_3CN$ containing 0.1 M TEAA) and eluent was desalted on a Sep-Pak C18 Cartridge (Waters) and evaporated under reduced pressure. The product was freeze-dried to afford **3** (68 mg, 0.17 mmol) as a pink solid, yield 15% (for 2 steps).

$^1$H-NMR (400 MHz, $CD_3OD$) δ 7.91 (d, *J* = 1.8 Hz, 1H), 7.60 (dd, *J* = 8.0, 2.1 Hz, 1H), 7.13 (d, *J* = 7.8 Hz, 1H), 6.62 (d, *J* = 8.2 Hz, 2H), 6.39 (d, *J* = 2.3 Hz, 2H), 6.29 (dd, *J* = 8.2, 2.3 Hz, 2H), 5.42 (s, 1H), 4.61 (s, 2H)

$^{13}$C-NMR (100 MHz, $CD_3OD$) δ 151.4, 147.5, 142.7, 138.8, 130.5, 129.8, 125.1, 124.7, 121.2, 113.2, 110.9, 102.0, 46.5, 7.9.

HRMS (ESI$^+$): *m/z* Calcd. for [M+H]$^+$, 399.10147, Found, 399.09779 (-3.68 mDa).

**Preparation of sHMRG (4)**

Leuco-sHMRG (10.0 mg, 0.025 mmol) and chloranil (18.5 mg, 0.075 mmol) were dissolved in *N*,*N*-dimethylformamide (1 mL) and stirred at RT for 10 min. Then, the mixture was purified by MPLC (eluent: A/B = 95/5 to 5/95, A: 100% $H_2O$ containing 0.1% TFA, B: 100% $CH_3CN$ containing 0.1% TFA) and eluent was freeze-dried to afford **4** (6.2 mg, 0.016 mmol), yield 62%.

$^1$H-NMR (400 MHz, DMSO-$d_6$) δ 7.95 (s, 1H), 7.65 (d, 1H, *J* = 8.4 Hz), 7.19 (d, 1H, *J* = 8.4 Hz), 6.98 (d, 2H, *J* = 8.8 Hz), 6.82 (d, 2H, *J* = 8.8 Hz), 6.73 (s, 2H), 4.12 (s, 2H).

$^{13}$C-NMR (100 MHz, DMSO-$d_6$) δ 160.1, 158.1, 157.4, 150.1, 140.5, 132.5, 130.5, 129.0, 125.5, 124.7, 117.5, 113.5, 97.6, 61.1.

HRMS (ESI$^+$): *m/z* Calcd. for [M+H]$^+$, 397.08582, Found, 397.08808 (+2.26 mDa).

**Preparation of sHMRG-based probes**

Peptide building blocks were prepared using the automated peptide synthesizer SyroI as ones protected with acid-labile protective groups for side chains and Fmoc group for *N*-terminal amino group.

Leuco-sHMRG (11 μmol), protected amino acids/peptides (3 μmol), and DMT-MM (5 μmol) were dissolved in *N*-methylpyrrolidone (NMP, 30 μL). The reaction was stirred at 40°C for 30 min. Chloranil (23 μmol) was added and stirred at RT for 10 min. The mixture was purified by MPLC (eluent: A/B = 95/5 to 5/95, A: 100% $H_2O$ containing 0.1% TFA, B: 100% $CH_3CN$ containing 0.1% TFA) and eluent was freeze-dried. If the building block had Fmoc group, piperidine (50 μL) and NMP (50 μL) were added to the residue and stirred at RT for 30 min. If the building block had acid-labile protective groups, TFA (1000 μL) was added to the residue or mixture and stirred at RT for 2 h. The mixture was purified by MPLC (eluent: A/B = 95/5 to 5/95, A: 100% $H_2O$ containing 0.1% TFA, B: 100% $CH_3CN$ containing 0.1% TFA) and eluent was freeze-dried to afford the product.

**Preparation of GP-sHMRG (5)**

GP-sHMRG was prepared using Fmoc-Gly-Pro-OH as the building block.

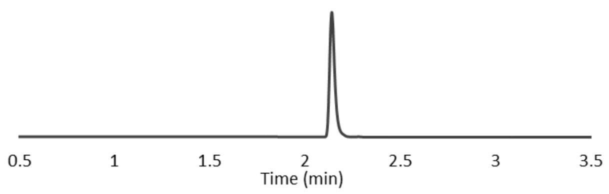


HRMS (ESI+): *m/z* Calcd. for [M+H]+, 551.16004, Found, 551.16154 (+1.50 mDa).

**Preparation of AcGP-sHMRG (6)**

Ac-GP-sHMRG was prepared using Ac-Gly-Pro-OH as the building block.

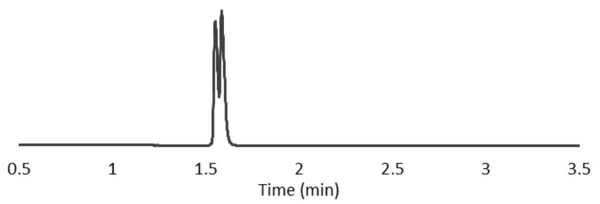


HRMS (ESI+): *m/z* Calcd. for [M+H]+, 593.17061, Found, 593.17082 (+0.21 mDa).

**Preparation of Y-sHMRG (7)**

Y-sHMRG was prepared using Boc-Tyr(*t*Bu)-OH as the building block.

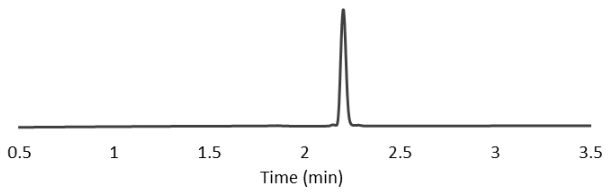


HRMS (ESI+): *m/z* Calcd. for [M+H]+, 560.14914, Found, 560.14900 (-0.14 mDa).

**Preparation of Ac-Met-sHMRG (8)**

Ac-Ala-sHMRG was prepared using Ac-Ala-OH as the building block.

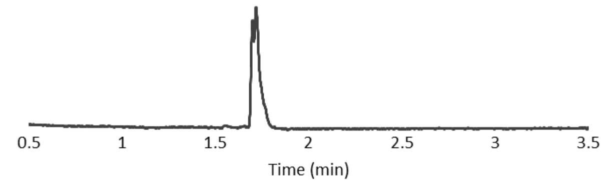


HRMS (ESI+): *m/z* Calcd. for $[M+Na]^+$, 532.11544, Found, 532.11747 (+2.03 mDa).

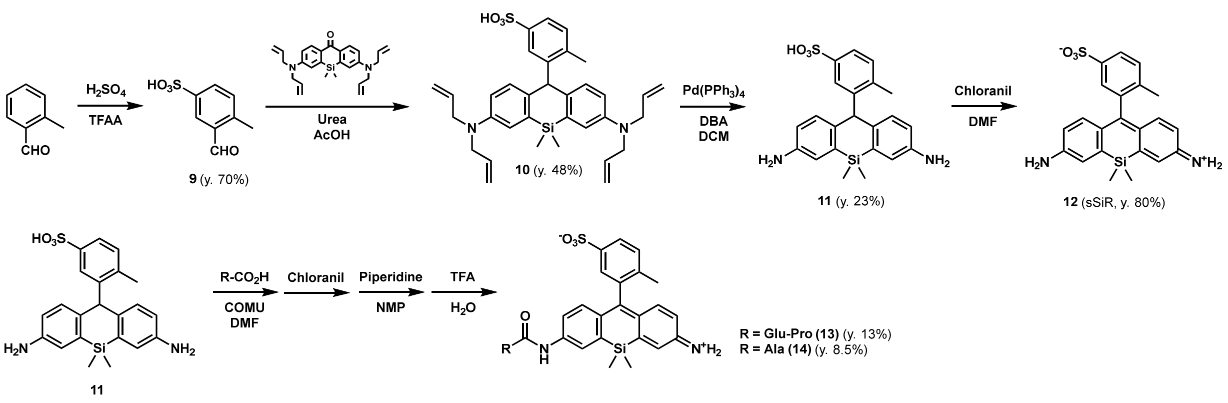


**Scheme S2.** Preparation of sSiR and fluorogenic probes for proteases/peptidases.

**Preparation of 3-formyl-4-methylbenzenesulfonic acid (9)**

Trifluoroacetic anhydride (TFAA; 6.0 mL, 43.7 mmol) was added to $H_2SO_4$ (1.1 mL, 20.8 mmol) with cooling at 0°C under Ar, then the solution was brought to room temperature and stirred for 3.5 h. 2-methylbenzaldehyde (2.5 mL, 20.8 mmol) was added dropwise under cooling at 0°C, then the solution was brought to room temperature and stirred for 20 h. Water was added dropwise under cooling at 0°C, and TFAA and water were removed in vacuo. The residue was purified by MPLC (eluent: A/B = 95/5 to 5/95, A: 100% $H_2O$ containing 0.1% TFA, B: 100% $CH_3CN$ containing 0.1% TFA) and eluent was evaporated and freeze-dried to afford **9** (2.9 g, 14.5 mmol) as a colorless solid, yield 70%.

$^1$H-NMR (400 MHz, $CD_3CN$) δ 10.27 (s, 1H), 8.22 (d, *J* = 1.8 Hz, 1H), 7.94 (dd, *J* = 8.0, 2.1 Hz, 1H), 7.50 (d, *J* = 7.8 Hz, 1H), 2.70 (s, 3H)

**Preparation of leuco-sSiR (11)**

*N*,*N*,*N'*,*N'*-tetraallyldiamino-Si-xanthone (1.77 g, 4.34 mmol), 3-formyl-4-methylbenzenesulfonic acid (868 mg, 4.34 mmol), and Urea (130 mg, 2.17 mmol) were dissolved in acetic acid (20 mL) and stirred at 90°C for 17 h under Ar, and then evaporated. The residue was purified by column chromatography on silica gel (eluent: dichloromethane/methanol) to afford **10** (1220 mg, 2.09 mmol), yield 48%.

**10** (350 mg, 0.616 mmol) and 1,3-dimethylbarbituric acid (577 mg, 3.70 mmol) were dissolved in dichloromethane (20 mL) and the solution was degassed using an aspirator. Then, tetrakis(triphenylphosphine)palladium(0) (28 mg, 0.024 mmol) was added, and the solvent was further degassed under reduced pressure. The reaction mixture was stirred at 35°C for 1 h under Ar, and then evaporated. The residue was purified by column chromatography (silica gel; dichloromethane/methanol). The eluent was evaporated to afford **11** (60 mg, 0.142 mmol), yield 23%.

$^1$H-NMR (400 MHz, $CD_3OD$) δ 7.60 (d, *J* = 1.8 Hz, 1H), 7.49 (dd, *J* = 8.2, 1.8 Hz, 1H), 7.15 (d, *J* = 8.2 Hz, 1H), 6.99 (d, *J* = 2.5 Hz, 2H), 6.79 (d, *J* = 8.5 Hz, 2H), 6.59 (dd, *J* = 8.5, 2.5 Hz, 2H), 5.59 (s, 1H), 2.25 (s, 3H), 0.58 (s, 3H), 0.36 (s, 3H)

$^{13}$C-NMR (100 MHz, $CD_3OD$) δ 148.0, 145.3, 143.1, 139.2, 138.7, 134.6, 131.9, 130.8, 129.1, 124.0, 120.4, 118.4, 51.1, 20.3, -0.4, -0.8

**Preparation of sSiR (12)**

Leuco-sSiR (211 mg, 0.497 mmol) and chloranil (246 mg, 1 mmol) were dissolved in *N*,*N*-dimethylformamide (4 mL) and dichloromethane (4 mL) and stirred at room temperature for 5 min. Then, the mixture was purified by column chromatography on silica gel (eluent: dichloromethane/methanol) to afford **12** (168 mg, 0.398 mmol), yield 80%.

$^1$H-NMR (400 MHz, $CD_3OD$) δ 7.88 (dd, *J* = 8.0, 1.8 Hz, 1H), 7.53 (d, *J* = 1.8 Hz, 1H), 7.46 (d, *J* = 8.0 Hz, 1H), 7.18 (d, *J* = 2.5 Hz, 2H), 6.97 (d, *J* = 9.4 Hz, 2H), 6.53 (dd, *J* = 9.4, 2.5 Hz, 2H), 2.05 (s, 3H), 0.50 (s, 3H), 0.49 (s, 3H)

$^{13}$C-NMR (100 MHz, $CD_3OD$) δ 168.8, 157.2, 149.0, 142.6, 142.3, 138.7, 138.1, 130.1, 126.9, 126.1, 123.3, 115.7, 109.5, 18.0, -2.8, -3.1

HRMS (ESI$^+$): *m/z* Calcd. for $[M+Na]^+$, 445.10181, Found, 445.10170 (-0.11 mDa).

**Preparation of sSiR-based probes**

Peptide building blocks were prepared using the automated peptide synthesizer SyroI as ones protected with acid-labile protective groups for side chains and Fmoc group for N-terminal amino group.

Leuco-sSiR (11.8 µmol), protected amino acids/peptides (4.7 µmol), and COMU (4.7 µmol) were dissolved in *N*,*N*-dimethylformamide (DMF, 60 µL). The reaction was stirred at room temperature for 30 min. Chloranil (23.5 µmol) was added and stirred at RT for 10 min. The mixture was purified by MPLC (eluent: A/B = 95/5 to 5/95 over 15 min, A: 100% $H_2O$ containing 0.1% TFA, B: 100% $CH_3CN$ containing 0.1% TFA) and eluent was freeze-dried. If the building block had Fmoc group, piperidine (50 µL) and NMP (50 µL) were added to the residue and stirred at RT for 30 min. If the building block had acid-labile protective groups, TFA (1000 µL) was added to the residue or mixture and stirred at RT for 2 h. The mixture was purified by MPLC (eluent: A/B = 95/5 to 5/95, A: 100% $H_2O$ containing 0.1% TFA, B: 100% $CH_3CN$ containing 0.1% TFA) and eluent was freeze-dried to obtain the product.

**Preparation of EP-sSiR (13)**

EP-sSiR was prepared using Fmoc-Glu(*t*Bu)-Pro-OH as the building block.

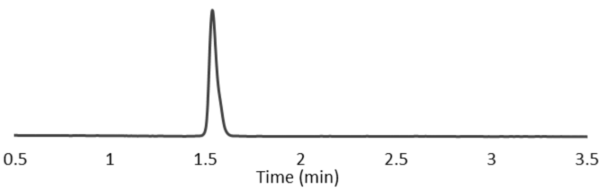


HRMS (ESI+): *m/z* Calcd. for [M+H]+, 649.21522, Found, 649.21491 (-0.31 mDa).

**Preparation of A-sSiR (14)**

A-sSiR was prepared using Boc-Ala-OH as the building block.

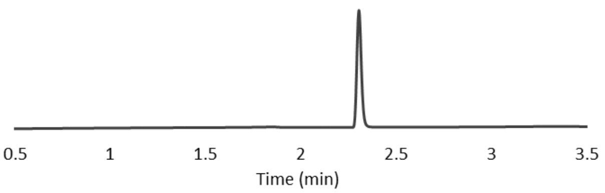


HRMS (ESI+): *m/z* Calcd. for [M+H]+, 494.15698, Found, 494.15693 (-0.05 mDa).

**Preparation of Ac-Ala-PMAC**

Ac-Ala-PMAC was prepared according to the procedure reported in the literature[9], using Ac-Ala-OH as a building block.

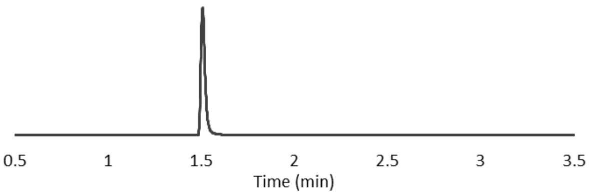


LRMS (ESI+): *m/z* = 369.1 [M+H]+.

### Supplementary References

[1] K. Honda, T. Okusaka, K. Felix, S. Nakamori, N. Sata, H. Nagai, T. Ioka, A. Tsuchida, T. Shimahara, M. Shimahara, Y. Yasunami, H. Kuwabara, T. Sakuma, Y. Otsuka, N. Ota, M. Shitashige, T. Kosuge, M. W. Büchler, T. Yamada, *PLoS One* **2012**, *7*, DOI 10.1371/journal.pone.0046908.

[2] K. Honda, M. Kobayashi, T. Okusaka, J. A. Rinaudo, Y. Huang, T. Marsh, M. Sanada, Y. Sasajima, S. Nakamori, M. Shimahara, T. Ueno, A. Tsuchida, N. Sata, T. Ioka, Y. Yasunami, T. Kosuge, N. Miura, M. Kamita, T. Sakamoto, H. Shoji, G. Jung, S. Srivastava, T. Yamada, *Sci. Rep.* **2015**, *5*, DOI 10.1038/srep15921.

[3] S. Kato, K. Honda, *Cancers (Basel)* **2020**, *12*, 1–21.

[4] K. Felix, K. Honda, K. Nagashima, A. Kashiro, K. Takeuchi, T. Kobayashi, S. Hinterkopf, M. M. Gaida, H. Dang, N. Brindl, J. Kaiser, M. W. Büchler, O. Strobel, *Int. J. Cancer* **2022**, *150*, 881–894.

[5] M. Sakabe, D. Asanuma, M. Kamiya, R. J. Iwatate, K. Hanaoka, T. Terai, T. Nagano, Y. Urano, *J. Am. Chem. Soc.* **2013**, *135*, 409–414.

[6] M. Minoda, J. Hatakeyama, N. Nagano, T. Mizuno, T. Iwasaka, S. Shiga, K. Takahashi, H. Hiraide, S. Sakamoto, Y. Kagami, A. Kashiro, K. Honda, Y. Sugiura, K. Mishima, M. K. Mishima, H. Kusuhara, Y. Urano, T. Komatsu, *J. Am. Chem. Soc.* **2025**, *147*, 4743–4751.

[7] H. Noji, Y. Minagawa, H. Ueno, *Lab Chip* **2022**, *22*, 3092–3109.

[8] T. Komatsu, T. Mizuno, *ACS Cent. Sci.* **2025**, *in press*, DOI 10.1021/acscentsci.5c00100.

[9] S. Sakamoto, H. Hiraide, M. Minoda, N. Iwakura, M. Suzuki, J. Ando, C. Takahashi, I. Takahashi, K. Murai, Y. Kagami, T. Mizuno, T. Koike, S. Nara, C. Morizane, S. Hijioka, A. Kashiro, K. Honda, R. Watanabe, Y. Urano, T. Komatsu, *Cell Rep. Methods* **2024**, *4*, 100688.

[10] T. Komatsu, K. Hanaoka, A. Adibekian, K. Yoshioka, T. Terai, T. Ueno, M. Kawaguchi, B. Cravatt, T. Nagano, *J. Am. Chem. Soc.* **2013**, *135*, 6002–6005.

[11] Y. Kushida, T. Nagano, K. Hanaoka, *Analyst* **2015**, *140*, 685–695.

[12] Y. Kushida, K. Hanaoka, T. Komatsu, T. Terai, T. Ueno, K. Yoshida, M. Uchiyama, T. Nagano, *Bioorg. Med. Chem. Lett.* **2012**, *22*, 3908–3911.
